# Cross-level correlation and temporal dynamics of hierarchical habitat selection

**DOI:** 10.64898/2026.11.13.764186

**Authors:** Léopold Berger, Charles P. Henriot, Thibaut Powolny, Régis Renaude, Mickaël Sage, Christophe Bonenfant

## Abstract

Habitat selection is a fundamentally hierarchical process implying that choices made by animals at broad spatial scales constrain those made at finer ones. How selection for the same habitats relates across hierarchical orders remains largely untested empirically. We investigated whether selection for a given habitat at the larger scale reinforces itself, is independent of, or opposes selection for the same habitat at the finer scale. Using GPS-collared European hares (*Lepus europaeus*, *n* = 52) in a mid-mountain grassland landscape (French Jura, 2021–2023), we jointly quantified and compared temporal variation in second-order (home range placement within the landscape) and third-order (habitat use within home ranges) selection with generalised linear mixed models, and resource selection functions. The association in selection strength between orders was strongly habitat-dependent: positive for grasslands, forests and farm tracks, and negative for fodder crops, although support for forests and fodder crops weakened when sex differences were accounted for. Our results suggest that hierarchical levels of habitat selection can couple or decouple depending on the habitat considered and what resources European hares are seeking at the different scales. Temporal variability in selection was consistently greater within home ranges than at the landscape level, and seasonal variation in ecological conditions (temperature, precipitation) shaped fine-scale selection while leaving landscape-level selection largely unaffected. Large-scale selection by European hare appeared relatively stable over time while fine-scale selection allowed dynamic behavioural adjustments to short-term variability of environmental conditions. Our results illustrate a hierarchical strategy of stable placement of the home range, flexible use within it. Reliance on heterogeneous open and semi-open habitats and consistent avoidance of built-up areas also clarifies the habitat requirements of the European hare in grassland-dominated agricultural landscapes, given population declines across much of its native range.

## Introduction

For mobile organisms, habitat selection is a fundamental determinant of fitness (Hildén 1965, Gaillard et al. 2010), shaping survival and reproductive success of individuals (McLoughlin et al. 2007), and results from behavioural decisions driven by resource availability, species interactions, and disturbance regimes (Roever et al. 2014, Desbiez et al. 2020, Beumer et al. 2023). Because individuals must trade-off competing needs such as foraging, reproduction, and predator avoidance (Mysterud and Ims 1998, Thomson et al. 2006), habitat selection is dynamic and its expression differs according to the scale of observation (Johnson 1980, Wiens 1989). Habitat selection has long been conceptualised as a hierarchical process, Johnson (1980) for instance, distinguishing four nested orders: first-order (geographical range), second-order (home range placement), third-order (habitat use within the home range), and fourth-order (fine-scale resource selection, *e.g.,* plant parts consumed by herbivores). Selection is now considered to operate at hierarchical levels across a continuum of spatial scales (McGarigal et al. 2016). Relationships between these levels can arise through top-down constraints, as broad-scale patterns of space use shape habitat availability at finer scales (Johnson 1980) and through bottom-up processes, as repeated fine-scale movement and habitat-selection decisions can generate broader patterns of space use (Picardi et al. 2024).

Most multi-level habitat selection studies on mammals and birds based on telemetry data focus on landscape and home range habitat selection (*e.g.,* Dupke et al. 2017, Quinlan et al. 2026) and empirical comparisons across hierarchical levels remain largely untested. Qualitative comparisons suggest that the relative ranking of habitats can differ among levels of habitat selection (raccoon *Procyon lotor*, Beasley et al. 2007; feral pig *Sus scrofa*, Paolini et al. 2019; roe deer *Capreolus capreolus*, Dupke et al. 2017; elk *Cervus canadensis*, Quinlan et al. 2026), likely because animals encounter habitats under different spatial configurations. Rare or spatially aggregated habitats at the landscape level may strongly shape home range placement, whereas widely distributed habitats may instead shape behavioural decisions within home ranges. Some studies have also compared the temporal dynamics of habitat selection across hierarchical levels (*e.g.,* Dupke et al. 2017), suggesting environmental conditions could generate temporal associations in selection strength akin to the climate forcing on population dynamics.

Spatial constraints on space use can also affect an animal’s ability to select a given habitat and decouple association between levels. Many studies report greater inter-seasonal or daily variation in habitat use at fine scales (*e.g.,* within home ranges), while seasonal ranges are more consistent at least on the short-term (Sarmento et al. 2006, Beatty et al. 2014, Dupke et al. 2017). In large herbivores, for instance, home range composition was more stable in time than finer-scale daily-area selection, which varied more across seasons (woodland caribou *Rangifer tarandus*, Rettie and Messier 2000; red deer *Cervus elaphus*, Fattebert et al. 2019). Major landscape-level changes (*e.g.,* floods, landslides) can nonetheless alter landscape selection between years, as when African buffalo (*Syncerus caffer*) shifted seasonal habitat use after a large flood (Bennitt et al. 2014). The degree of philopatry or territoriality may, however, modulate this temporal stability as exemplified by the strongly philopatric roe deer (Wahlström and Liberg 1995) that showed little home range relocation after the 1999 Lothar storm despite major changes in food availability and vegetation structure (Widmer et al. 2004). Landscape-level habitat selection is therefore generally more stable in time than habitat selection within home ranges.

The relative spatio-temporal variability in landscape- and home range-level habitat selection behaviour matches the idea that coarse-scale selection reflects long-term limiting factors while fine-scale selection responds to more immediate factors and needs (Rettie and Messier 2000). Animals often select home range locations to secure long-term access to key resources, hence anticipating predictable seasonal needs (*e.g.,* Shakeri et al. (2021) on mountain goat, *Oreamnos americanus*). Within home range selection is frequently responsive to short-term variation in forage, predation risk, or human activity. Because home range characteristics such as size and composition are related to habitat selection and use of space within the range (Johnson 1980), the relative constraints in habitat selection behaviour at the different levels may generate temporal relationships in the strength of selection over time. In feral pigs, both resource selection and home range size varied seasonally, leading Paolini et al. (2019) to suggest that second- and third-order selection were mechanistically linked through a habitat “functional response”. However, the current lack of empirical and theoretical studies makes it difficult to predict the direction of this cross-level relationship at the population level (see Van Moorter et al. 2016, for an attempt at the individual level), as habitat selection has mostly been studied at each hierarchical level separately rather than by explicitly relating them through time.

The European hare (*Lepus europaeus*) is historically associated with human agricultural practices and typically inhabits heterogeneous open farmland landscapes, where individuals maintain relatively stable home ranges (Hackländer 2022) once established (Bray et al. 2007), making the species a useful model for examining habitat selection across hierarchical levels. Most habitat selection studies on this species have focused on habitat selection within the home range (home range-level selection) in lowland agricultural landscapes (*e.g.,* Petrovan et al. 2013, Ullmann et al. 2020), whereas fewer studies have examined how individuals position their home ranges within the broader landscape (landscape-level selection). Consequently, explicit comparisons of habitat selection across these two hierarchical levels remain lacking. Using GPS-collared European hares from a reinforcement programme in France that exhibited stable seasonal home ranges (*n* = 52), we investigated habitat selection at two hierarchical levels: home range placement within the landscape (second-order selection) and habitat selection within home ranges (third-order selection), treating the relationship between levels as a biological response variable. Our objectives were to (1) quantify temporal variation in habitat selection at these two hierarchical levels; (2) quantify the strength and direction of temporal relationships in habitat selection between the second and third orders; and (3) assess how temporal environmental variables influence habitat selection at each order and may contribute to relationships between them.

A first hypothesis linking selection across scales is that stronger selection for a habitat at the landscape level should result in greater availability of that habitat within the home range, leading to weaker selection for that same habitat at the home range level. Accordingly we predicted (H1) that habitat selection of European hares would be negatively related between the two levels for all habitat types. A second hypothesis is that individuals faithful to their home ranges have little scope for changes in second-order selection, while remaining more flexible in their habitat use within established home ranges. European hares are known for their high spatial fidelity and relatively limited shifts in home range location (Rühe and Hohmann 2004, Bray et al. 2007, Hackländer 2022). We hence predicted (H2) a decreasing correlation in habitat selection between landscape and home range scales with the degree of site fidelity of hares. Finally, short-term variation in weather (temperature, precipitation) can alter life conditions of hares and their resource availability. In response, individuals can adjust habitat use within an established home range without necessarily relocating it (Mayer et al. 2019, Hackländer 2022). Accordingly, we predicted (H3) that short-term variation in environmental conditions would influence habitat selection in the same direction at both the landscape and home range levels, hence generating a positive association between selection levels. The strength of this effect should however be greater and its response more immediate at the home range than the landscape level because fine-scale movements within the home range allow individuals to respond more directly to short-term changing conditions.

## Materials and methods

### Biological model

The European hare is a medium-sized, solitary, and non-territorial herbivore adapted to open cropland and grassland habitats (Smith et al. 2005, Hackländer 2022, Mori et al. 2022). Its diet, mainly grasses and herbaceous plants, varies seasonally with vegetation availability and nutritional quality (Reichlin et al. 2006, Mori et al. 2022, Santilli et al. 2024), and its high energetic and protein requirements, especially during reproduction, make it sensitive to changes in food resources and habitat structure (Schai-Braun et al. 2015). Following translocation, wild hares typically settle within a few weeks to two months (Fischer and Tagand 2012). Hares rest in shallow forms that are regularly relocated and selected for concealment, escape opportunities, and access to foraging areas (Neumann et al. 2012).

### Study area

The study area (50 km^2^) is located in the Jura Mountains of eastern France (46.98°–47.06° N, 6.00°–6.12° E; WGS84) and consists of a grassland-dominated landscape ranging from 350 to 860 m a.s.l. The area is characterised by a hilly topography and a heterogeneous habitat mosaic, comprising 43% grasslands, 38% forests (including 15% forest edges), 7% arable land, 4% woodlands and hedgerows, 4% built-up areas, 2% roads, and 2% other open fields. Grasslands are predominantly permanent and managed either for extensive cattle grazing or hay production within a dairy farming system dedicated to Comté cheese production. These grasslands are mainly represented by the *Heracleo sphondylii–Brometum mollis* association (B. Foucault 1989 ex 2008) and the *Lolio perennis–Cynosuretum cristati* association (Braun-Blanq. & de Leeuw, Tüxen 1937). The climate is semi-continental, with a mean annual rainfall of 1233 mm and a mean annual temperature of 11.1 °C during the 2021–2023 survival period of released hares. Mean seasonal rainfalls (2021–2023) were 305, 288, 415, and 226 mm in spring, summer, autumn, and winter, respectively, while mean seasonal temperatures were 13.6, 18.5, 7.8, and 4.5 °C for the same seasons. Winter conditions show intermittent snow cover from November to April, with snow depths of around 5–10 cm.

Our study area hosts a guild of mid-size mammalian predators that prey upon European hares, with the red fox (*Vulpes vulpes*) identified as the primary predator and mustelids (*e.g.,* European pine marten (*Martes martes*), stoat (*Mustela erminea*)) as secondary predators (Henriot et al. 2025). Other potential predators present in the study area include raptors, wildcats (*Felis silvestris*), and lynx (*Lynx lynx*). Although the European hare is a game species in France including the study area, hunting of this species was suspended in the study area from 2019 to 2023, *i.e.,* two years before and throughout the GPS-tracking period (2021–2023), to minimise the loss of tagged individuals. Spring surveys using distance sampling indicate relatively stable and low hare abundance in the study area, ranging from 2.7 to 4.1 hares·km^-1^ between 2019 and 2023 (Henriot et al. 2025).

### Rearing and GPS monitoring

We conducted this study over a 2-year period from July 2021 to June 2023. All monitored European hares were captive-bred at a licensed farm (GAEC Tarral, Aveyron, France) operating under national regulations (French Environment Code, articles L413-1, L413-5) and underwent at least one month of acclimatisation in outdoor enclosures before release (see Henriot et al. 2025). We released a total of *n* = 103 subadult hares (5–6 months old; mean body mass = _3.87_ 3.92 _3.97_ kg; sex ratio 1:1) at six different sites in summer 2021 and summer 2022. Before release in the wild, we equipped each hare with a GPS collar (BioLoc Tx, XERIUS, Toulouse, France) integrating a UHF transmitter and a tri-axial accelerometer. Collars weighed 55 g, corresponding to <2% of body mass, well below recommended thresholds (Sikes et al. 2016). Devices were specifically designed and field-tested for the European hare prior to release. Handling time was <2 min per individual, without anaesthesia and in a quiet environment to minimise stress.

We programmed the GPS devices to record one location per hour for 24 h, followed by one location every 6 h (00:00, 06:00, 12:00, 18:00 GMT0) for the next 5 days. This 6-day acquisition cycle was repeated throughout the tracking period, ensuring that both foraging/active and resting/inactive behavioural states were sampled (Schai-Braun et al. 2012). This recording schedule would have theoretically allowed continuous monitoring for up to one year. Location record was carried out until the battery ran out or the animal died (more details in Henriot et al. 2025). Aside from a few clear outliers, we retained all GPS locations because position accuracy could not be reliably filtered using conventional quality metrics (*e.g.,* HDOP, number of satellites), which are often weak predictors of true location error in comparable GPS collar systems (Fleming et al. 2020).

### Data preparation

#### Seasonal home range estimation

Home range estimators require stationarity of locations to be meaningful (Péron 2019). To ensure that locations were stationary, we excluded GPS points recorded immediately after release for all individuals until a stable movement pattern was observed (Morellet et al. 2009, Mayer et al. 2021). We identified stable movement visually on QGIS following Péron (2019), Morellet et al. (2009), and Mayer et al. (2021). In a second step, we classified seasonal home ranges as stationary or non-stationary based on empirical variogram analyses and retained only stationary seasonal home ranges for subsequent analyses (see Atmeh et al. 2024). In addition, we included only seasonal home ranges estimated from stationarity periods representing at least 20% (18 days) of the corresponding seasonal duration. This threshold ensured that stationarity reflected a meaningful portion of the season (100% ∼ either 90 or 91 days), rather than short-term residency or transient movements.

We estimated seasonal home ranges from GPS data with the weighted autocorrelated kernel density estimator (wAKDE) implemented in the *ctmm* R package (Calabrese et al. 2016) to account for irregular sampling intervals. We distinguished four seasons by seeking the best compromise between capturing temporal variability and maintaining sufficient data within each period. We divided the year into four three-month periods corresponding to key stages of the European hare’s reproductive cycle and (*e.g.,* Jennings et al. 2006) to major agricultural activities: winter (January–March; onset of reproduction), spring (April–June; reproductive peak and mowing), summer (July–September; harvest and continued mowing), and autumn (October–December; post-harvest, sowing, and non-reproductive period).

#### Home range fidelity

We assessed seasonal home range fidelity within individuals using the Sørensen overlap index (SI) (Richard et al. 2014), calculated from home range polygons, and the Bhattacharyya coefficient (BC), derived from utilisation distributions (Winner et al. 2018), computed between the seasonal home ranges of each individual. The SI quantifies the consistency of home range placement (second-order selection), whereas the BC also accounts for intensity of use.

#### Environmental covariates

We retrieved land-cover classes from the *Registre Parcellaire Graphique* (RPG, 2021–2023; IGN), the *IGN TOPO* vector dataset (2022), and *OpenStreetMap* (OSM) layers. We grouped and processed them into 10 ecologically relevant habitats that potentially influence European hare habitat selection (Hackländer 2022): grasslands, winter cereals, fodder crops, other open fields (*e.g.,* maize and winter rapeseed), forests, forest edges, woodlands and hedgerows, farm tracks, secondary roads, and built-up areas (Supporting information). All spatial data were projected in EPSG:2154 (Lambert-93) and processed at a 5 m spatial resolution, providing an appropriate compromise between GPS positional accuracy and the spatial detail required to quantify fine-scale landscape features such as hedgerows. We created one raster map from land-cover layers for each year (2021, 2022, and 2023). Note that we used the 2021 map for summer 2021, the 2022 map for autumn 2021 to summer 2022, and the 2023 map for autumn 2022 to spring 2023, ensuring temporal consistency between habitat data and animal tracking periods. We conducted all spatial processing and map generation using QGIS 3.40.12 (QGIS.org 2026).

We characterised environmental conditions using mean seasonal temperature (°C) and mean seasonal precipitation (mm day^-1^), averaged from daily records of the *Météo-France* station #25223002 (Coulans), 2.5 km from the study area, and mean seasonal NDVI (Pettorelli et al. 2005), extracted from *Sentinel-2* imagery (10 m resolution) via the *Copernicus Browser* and used as a proxy for vegetation productivity.

### Habitat selection analyses

#### Landscape habitat selection (2^nd^ order)

We quantified the habitat selection following Johnson (1980) by estimating selection ratios (SRs) using the *adehabitatHS* R package (Calenge 2011). We defined habitat availability as identical for all individuals within a functionally accessible landscape (*A_L_*), delineated using terrain and land-cover data, and informed by the spatial extent of all available GPS locations. For each individual, we intersected seasonal home ranges with a land-cover map to calculate the proportional area of each of the 10 habitat types, *i.e.,* the used habitat (*U_i,t_*). We then calculated SRs as the ratio of the proportion of used habitat to its availability for each habitat type (*U_i,t_*/*A_L_*), where SR values >1 indicate selection, <1 indicate avoidance, and close to 1 values indicate proportional use (Manly et al. 2002). As a consequence, SR = 0 when a habitat type was available within *A_L_* but unused within an individual’s seasonal home range.

We analysed SRs using generalised linear mixed models (GLMMs) with a compound Poisson-Gamma Tweedie error distribution (1 < *ξ* < 2), which accommodates continuous non-negative responses with exact zeros (Hasan and Dunn 2011); its variance is a power function of the mean (Var(Y) = *φμ^ξ^*), with dispersion parameter *φ* and index parameter *ξ* estimated from the data. Fixed effects included habitat type (with grassland, the dominant available habitat, as the reference level), season-year (an eight-level categorical variable from summer 2021 to spring 2023), sex, and time since release (days since release at the midpoint of each seasonal tracking period), the latter as a nuisance variable accounting for post-release acclimatisation and unequal individual contributions across seasons arising from differential survival. We included two-way interactions between habitat type and sex, because we expected males and females to behave differently, and between habitat type and season-year, because temporal variation in selection may differ among habitats. Hare identity was entered as a random intercept (Bolker et al. 2009).

#### Home range habitat selection (3^rd^ order)

We quantified home range habitat selection using resource selection functions (RSFs; Manly et al. 2002), randomly sampling available points within each seasonal home range at a 1:10 use-available ratio (*e.g.,* Beers and Frey 2022). We fitted binomial GLMMs with a logit link and the same fixed effects and interactions as the landscape-level models, with individual random intercepts and uncorrelated random slopes for each habitat type (Muff et al. 2020) (Supporting information). We excluded built-up areas because their availability was <1% across all home ranges. We exponentiated model coefficients (*β*) into odds ratios (ORs), expressed relative to the sex-specific grand mean across all habitat types and season-years, so that values >1 indicate above-average and values <1 below-average relative selection. We compared the resulting habitat-, sex- and season-year-specific coefficients with those of the landscape level to assess H1.

#### Habitat selection and temporal environmental covariates

To evaluate H3, we replaced season-year in the landscape- and home range-level models with environmental covariates (temperature, precipitation, and NDVI), entered either individually or in pairs representing distinct abiotic drivers (*e.g.,* thermal conditions versus moisture-productivity gradients). We never combined NDVI and precipitation, which were strongly correlated (|*r* | > 0.7; Dormann et al. 2013). We quantified the proportion of the temporal variation in mean SRs and RSF coefficients (captured by season-year) explained by each covariate with the R^2^_ANODEV_ statistic (Skalski 1996), and computed population-average marginal slopes (*emtrends*) and predictions (*emmeans*) across the observed covariate range.

#### Temporal correlation between landscape and home range habitat selection

To test the association between habitat selection at the landscape and home range levels across the eight monitoring seasons, we quantified the relationship between the corresponding model-derived selection coefficients. Because both landscape-level SRs and home range-level RSF coefficients are estimated quantities subject to sampling uncertainty, we used an error-in-variables (EIV) linear regression (Fuller 1987), or Deming regression, which explicitly accounts for measurement error in both response and predictor variables (Linnet 1993), yielding a slope (*s*) that quantifies the strength and direction of coupling between selection at both orders. The measurement-error variance ratio *λ* was estimated as the ratio between the mean squared standard errors of the marginal means from the RSF coefficients (home range-level) and the SRs (landscape-level) derived from the temporal models. We conducted a global analysis pooling all habitat types to assess the overall correspondence between the landscape and home range levels of selection. We then fitted habitat-specific EIV regressions to evaluate whether the strength of the association between landscape and home range habitat selection differed among habitat types. Because EIV models rely on assumptions about the relative magnitude of measurement error in the two variables, we additionally fitted standard linear regressions as a sensitivity analysis (Supporting information). Models were fitted using the *SimplyAgree* R package (Caldwell 2022).

#### Site fidelity and cross-level coupling

To test H2, we analysed the change in habitat use and availability between two consecutive seasonal home ranges (called transition), and measured its fidelity with the Bhattacharyya coefficient. For each transition and habitat type, we computed the change in the log proportion of the habitat among available points (Δ log *A*, difference in availability within the home range between seasons *j* and *j* + 1) and among GPS locations (Δ log *U*, difference in use between seasons *j* and *j* + 1). Because a habitat may have no location in a given season, we added 0.5 to each count and 1 to the total before taking logarithms (Haldane 1956). We computed the binomial sampling variances of log proportions with the delta method. We retained habitat types with at least 20 available points in both home ranges. Within a land-cover map year, availability at the landscape level does not change, so that Δ log *A* equals the change in landscape-level selection, while the change in home range-level selection is Δ log *U* − Δ log *A*. We therefore modelled the change in use, Δ log *U*, with the change in availability, Δ log *A*, entered as an offset. The change in home range-level selection and its slope on Δ log *A* is therefore the cross-level coupling effect *c*. When *c* = 0, use changes in proportion to availability and selection is not coupled across levels; when *c* = −1, habitat availability does not affect its use.

To estimate *c*, we fitted linear mixed models considering Δ log *U* as the response variable, Δ log *A* as an offset, and Δ log *A*, centred fidelity and the two-way interaction as explanatory variables. We added a random slope of Δ log *A* for each transition because the hares’ response to changes in availability might vary among transitions. The model was of the form:

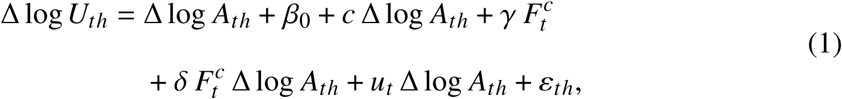

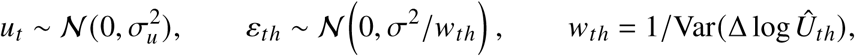

where *t* and ℎ index transitions and habitat types, *β*_0_ is the intercept, *γ* the main effect of fidelity, 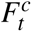 is the centred fidelity of transition *t*, *u_t_* is the transition-specific deviation of the coupling, and each observation of the response variable is weighted by the inverse of its sampling variance (*w_t_*_ℎ_). Under this model specification, H2 predicts a positive interaction (*δ* > 0), that is, a cross-level coupling closer to zero at higher fidelity.

We tested the interaction with a parametric bootstrap likelihood-ratio test (1,000 simulations; *pbkrtest* R package, Halekoh and Højsgaard 2014) and with a two-stage approach. We first regressed transition-specific couplings, estimated by weighted least squares within each transition, on fidelity, weighting each coupling by the inverse of its sampling variance and, as a check, without weights. Because fidelity varies only among transitions, the effective sample size for the interaction is the number of transitions rather than the number of transition × habitat observations. The main analysis used transitions within a land-cover map year, so that crop rotation could not change availability. We assessed if this restriction did drive our results by repeating the analysis with all consecutive transitions, including those spanning two land-cover maps for which availability may also change through crop rotation. We also assessed the influence of single transitions by refitting the two-stage regression after removing each transition in turn (leave-one-out): a transition was considered influential when its removal changed the conclusion of the test (*p* crossing 0.05). Finally, we replicated the same analysis by replacing fidelity with each individual’s centred CV of the availability of each habitat type across its seasonal home ranges, a habitat-specific measure of the stability of home range composition, and by adding a habitat-specific random slope of Δ log *A*. Because a higher CV indicates lower fidelity, H2 then predicts a negative interaction in this case. We fitted all linear mixed models using the *lme4* R package (Bates et al. 2015). A complementary habitat-level analysis is presented in Supporting information.

#### Model selection and diagnostics

We standardised continuous predictors (*z*-scores) to facilitate model convergence and selected models using the Bayesian Information Criterion (BIC; Burnham and Anderson 2002), retaining the model with the lowest BIC as the most parsimonious. We report point estimates with their 95% confidence intervals. We fitted all GLMMs with the *glmmTMB* R package (Brooks et al. 2017) and checked dispersion, zero inflation, and residual structure with the *DHARMa* R package (Hartig 2024), as well as random-effect singularity and multicollinearity with the *performance* R package (Lüdecke et al. 2021). We computed estimated marginal means and contrasts with the *emmeans* R package (Lenth and Piaskowski 2025) and back-transformed them to the original scale. We checked Deming regression residuals graphically for normality and homogeneity of variance. We conducted all analyses in R (version 4.4.1; R Core Team 2024).

#### Sample size

The sample size was determined by the reinforcement programme: all 103 hares released during the two sessions were GPS-collared and monitored. We retained every hare with at least one stationary seasonal home range covering at least 20% of a season, because home ranges and habitat selection within them can only be defined for resident animals. Because this sample could not be increased, we evaluated it through the precision of the estimates (Lakens 2022), and report 95% confidence intervals throughout. By design the cross-level slope comparison rests on 16 season-year-by-sex estimates per habitat type, and the interaction between availability change and fidelity on 33 transitions within an annual land-cover map (48 overall). Note that because the difference in habitat selection patterns between sexes was additive, our analysis of cross-level slopes mostly relies on the information of eight seasons.

## Results

### Fate of the released hares

Of the 103 hares released, 42 died or were lost to monitoring before establishing a seasonal home range, 40 of them within 14 days of release (Henriot et al. 2025), three were not monitored for long enough to cover 20% of a season, and six were excluded *a priori* because their locations were non-stationary throughout the entire study period. This gave a final sample of *n* = 52 hares, balanced between the sexes (26 females, 26 males), and 108 seasonal home ranges, with 3 to 12 hares per combination of season-year and sex. We carried out habitat selection analyses on 50,606 GPS locations from these 52 hares. Among them, 22 individuals (42%) were stationary in all seasons in which they were monitored (11 of them over a single season), whereas 30 individuals (58%) were stationary in some seasons only. The monitored hares had a mean 95% home range of _38.3_ 52.4 _66.6_ ha as estimated using the wAKDE method (Supporting information). Males exhibited substantially larger home ranges (_50.9_ 76.2 _101.4_ ha) than females (_23.0_ 28.7 _34.5_ ha) (*t*-test, *t* = −3.77, *df* = 27.59, *p* < 0.001).

### Habitat selection at the landscape level (second-order)

At the landscape level, a model estimating habitat selection of hares for each season-year (Fig. 1a) showed some temporal variation and differences between sexes. Across seasons, males and females selected woodlands and hedgerows, fodder crops, winter cereals, and grasslands, with the strongest selection for winter cereals (_1.88_ 2.72 _3.93_) and fodder crops (_1.38_ 2.36 _4.03_) in females, and for fodder crops (_1.25_ 2.03 _3.31_) and woodlands and hedgerows (_1.43_ 1.65 _1.90_) in males. Both sexes avoided forests and built-up areas. The sexes differed mainly for other open fields, selected by females (_1.14_ 1.44 _1.81_) but avoided by males (_0.54_ 0.69 _0.89_), for secondary roads, selected by females only (_1.28_ 1.86 _2.70_), and for forest edges, avoided by males only (_0.61_ 0.71 _0.83_). Farm tracks were used in proportion to availability by both sexes. According to the BIC ranking, the selected model however included time since release and the interaction between habitat type and sex only (Table 1a, Supporting information). The patterns of habitat selection hence differed between sexes only in their mean level of selection for each habitat, and seasonal variation was weak (Table 1a). Although the interaction between habitat type and season-year was detectable (Wald test, *χ*^2^ = 103.6, *df* = 63, *p* = 0.001), it was heavily penalised by the BIC. The largest seasonal fluctuations were observed for fodder crops, forests and winter cereals (Fig. 1a).

**Figure 1.**
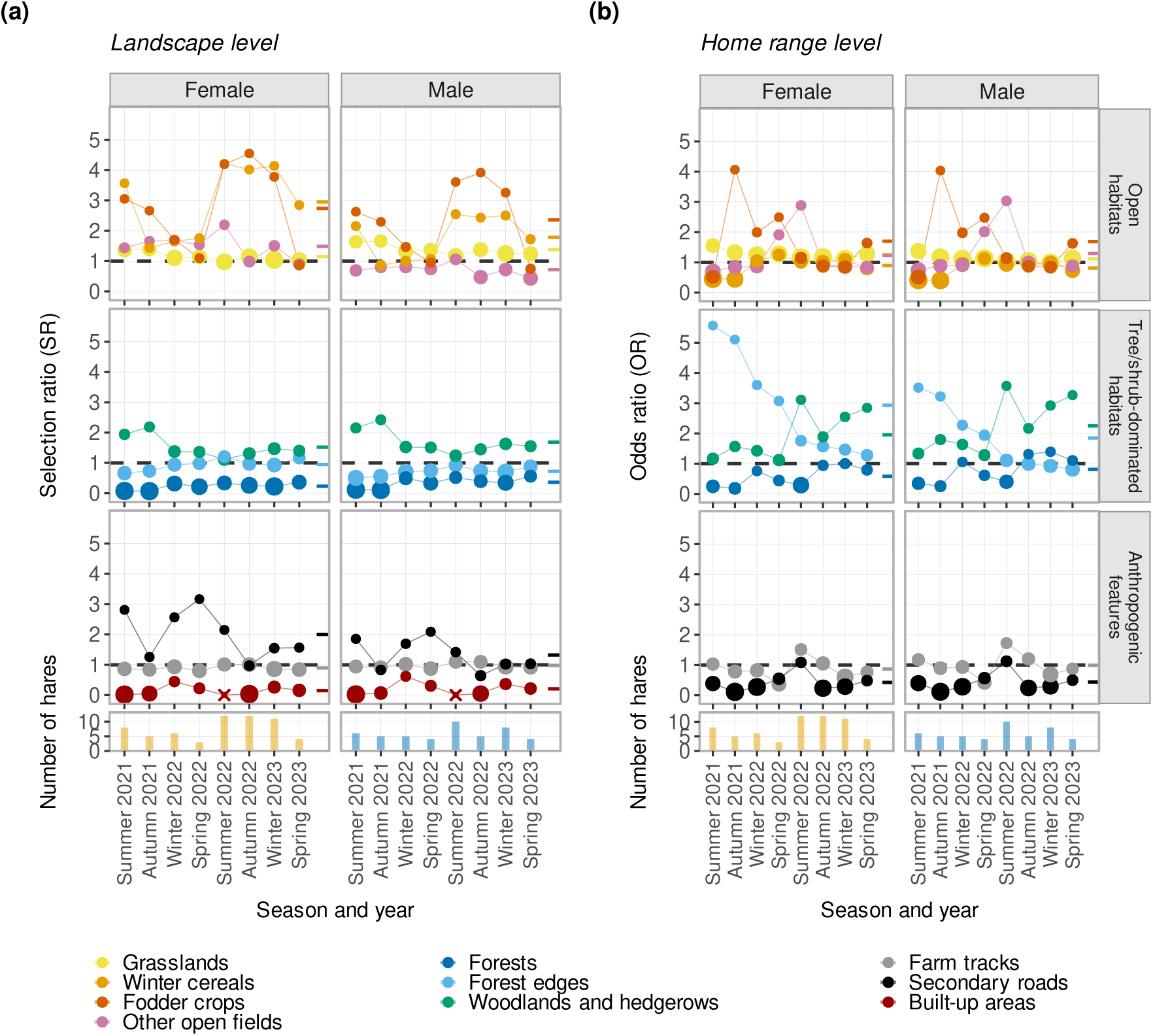
(a) Landscape and (b) home range habitat selection of released European hare (*Lepus europaeus*), by sex, across land-cover classes, from summer 2021 to spring 2023 in the Jura Mountains, France (52 hares, 108 seasonal home ranges). Habitat selection is quantified using predicted selection ratios (SRs) for landscape-level analyses and odds ratios (ORs) for home range-level analyses. Habitats with SRs or ORs > 1 are considered selected, whereas those with values < 1 are considered avoided. Mean values of SRs and ORs by habitat type are displayed as coloured dashes on the right side of each plot. Built-up areas were excluded from the home range-level analysis because they represented less than 1% of the individual-specific availability domain. Point size reflects estimate precision (inverse of the 95% confidence interval (CI) width). The crosses represent predictions for which CI could not be calculated due to high variability and proximity to 0. Histograms at the bottom of the panels show the number of hares available per season for habitat selection analyses, by sex.

**Table 1.**
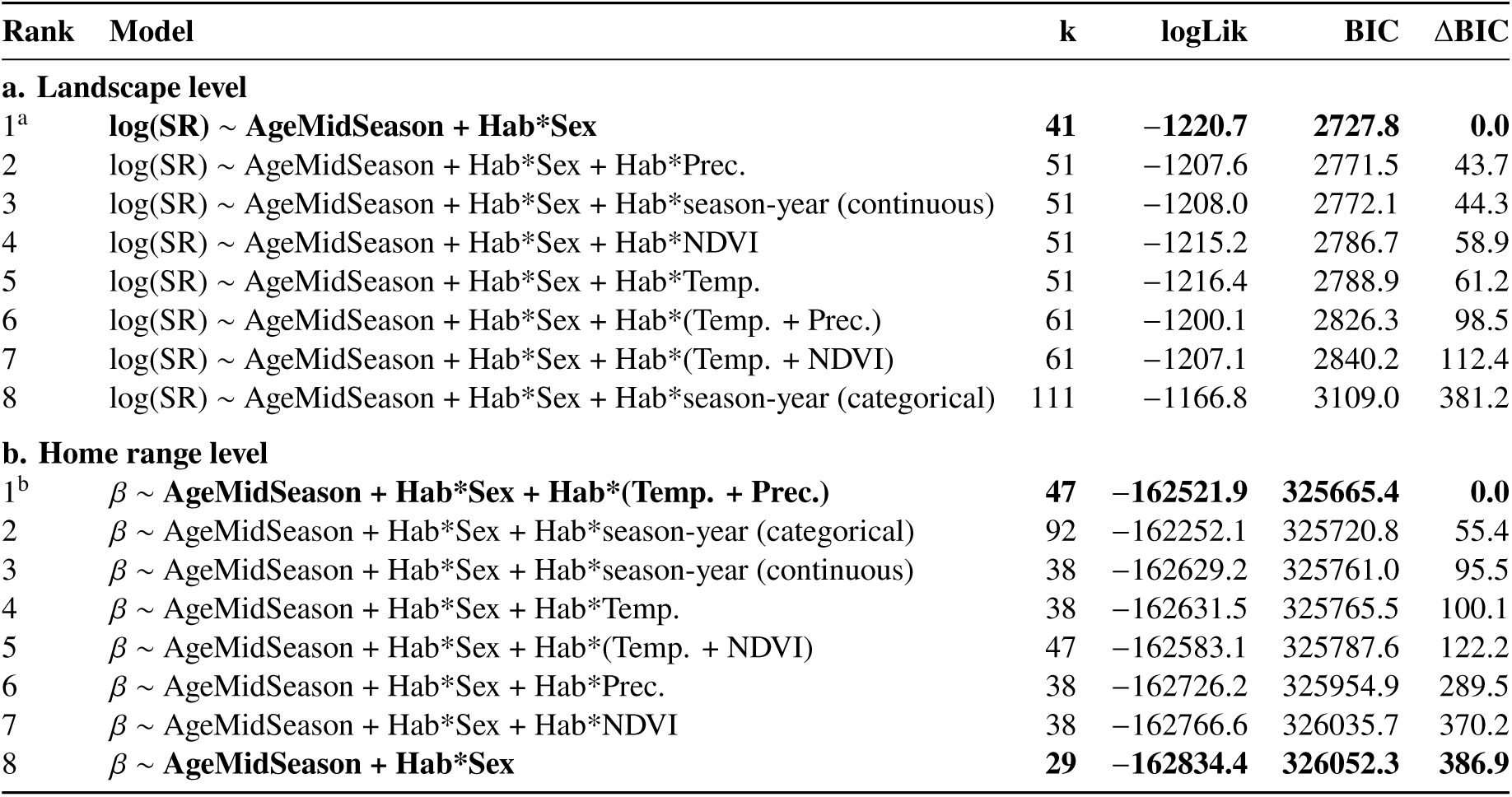
Comparison of the different models exploring habitat selection at (a) the landscape level and (b) the home range level of European hares (*Lepus europaeus*) released in a mid-mountain grassland landscape in the Jura Mountains, France, from summer 2021 to spring 2023 (52 hares, 108 seasonal home ranges). log(SR): landscape-level selection (ratio) coefficient. *β*: home range-level selection coefficient. AgeMidSeason: time since release (days since release at the midpoint of each seasonal tracking period). Hab: habitat. Season-year was considered either as an ordered covariate (testing trends) or as a categorical variable (testing temporal variation). Environmental covariates: Temp.: temperature; Prec.: precipitation; NDVI: normalised difference vegetation index. Degrees of freedom (k), log-likelihood (logLik), Bayesian information criterion (BIC), and ΔBIC values are provided. The constant model is in bold. a and b indicate the selected models at the landscape and home range levels, respectively.

| Rank | Model | k | logLik | BIC | ΔBIC |
|---|---|---|---|---|---|
| **a. Landscape level** |  |  |  |  |  |
| 1<sup>a</sup> | **log(SR) ~ AgeMidSeason + Hab\*Sex** | **41** | **−1220.7** | **2727.8** | **0.0** |
| 2 | log(SR) ~ AgeMidSeason + Hab*Sex + Hab*Prec. | 51 | −1207.6 | 2771.5 | 43.7 |
| 3 | log(SR) ~ AgeMidSeason + Hab*Sex + Hab*season-year (continuous) | 51 | −1208.0 | 2772.1 | 44.3 |
| 4 | log(SR) ~ AgeMidSeason + Hab*Sex + Hab*NDVI | 51 | −1215.2 | 2786.7 | 58.9 |
| 5 | log(SR) ~ AgeMidSeason + Hab*Sex + Hab*Temp. | 51 | −1216.4 | 2788.9 | 61.2 |
| 6 | log(SR) ~ AgeMidSeason + Hab*Sex + Hab*(Temp. + Prec.) | 61 | −1200.1 | 2826.3 | 98.5 |
| 7 | log(SR) ~ AgeMidSeason + Hab*Sex + Hab*(Temp. + NDVI) | 61 | −1207.1 | 2840.2 | 112.4 |
| 8 | log(SR) ~ AgeMidSeason + Hab*Sex + Hab*season-year (categorical) | 111 | −1166.8 | 3109.0 | 381.2 |
| **b. Home range level** |  |  |  |  |  |
| 1<sup>b</sup> | *β* ~ **AgeMidSeason + Hab\*Sex + Hab\*(Temp. + Prec.)** | **47** | **−162521.9** | **325665.4** | **0.0** |
| 2 | *β* ~ AgeMidSeason + Hab\*Sex + Hab\*season-year (categorical) | 92 | −162252.1 | 325720.8 | 55.4 |
| 3 | *β* ~ AgeMidSeason + Hab\*Sex + Hab\*season-year (continuous) | 38 | −162629.2 | 325761.0 | 95.5 |
| 4 | *β* ~ AgeMidSeason + Hab\*Sex + Hab\*Temp. | 38 | −162631.5 | 325765.5 | 100.1 |
| 5 | *β* ~ AgeMidSeason + Hab\*Sex + Hab\*(Temp. + NDVI) | 47 | −162583.1 | 325787.6 | 122.2 |
| 6 | *β* ~ AgeMidSeason + Hab\*Sex + Hab\*Prec. | 38 | −162726.2 | 325954.9 | 289.5 |
| 7 | *β* ~ AgeMidSeason + Hab\*Sex + Hab\*NDVI | 38 | −162766.6 | 326035.7 | 370.2 |
| 8 | *β* ~ **AgeMidSeason + Hab\*Sex** | **29** | **−162834.4** | **326052.3** | **386.9** |

### Habitat selection at the home range level (third-order)

At the home range level, the model best describing our data (according to the BIC) included the interactions between habitat type and both temperature and precipitation (Table 1b, Supporting information). We described seasonal patterns using the same model structure as at the landscape level (Fig. 1b; all estimates in Supporting information). The interaction between habitat type and season-year was strongly supported at this level (Wald test, *χ*^2^ = 1047.6, *df* = 56, *p* < 0.001). Both sexes selected cover habitats: females selected forest edges (_1.48_ 2.52 _4.29_) and males woodlands and hedgerows (_1.16_ 2.09 _3.79_), each habitat showing a tendency towards selection in the other sex. Both sexes clearly avoided secondary roads (females: _0.18_ 0.35 _0.68_; males: _0.19_ 0.36 _0.67_). No other habitat type was clearly selected or avoided.

### Cross-order coupling of habitat selection

Our prediction of a negative association between habitat selection at the landscape and home range levels (H1) was supported for only one habitat type (Fig. 2, Table 2, Supporting information). When pooling across habitats, the overall association between landscape and home range selection was positive though not significant (*s* = _−0.85_ 1.37 _3.58_). The observed associations varied among habitats, being positive for 3 of the 9 habitat types (33.3%: farm tracks, forests, and grasslands), negative for 1 (11.1%: fodder crops), and non-significant for 5 (55.6%) (Table 2). Secondary roads exhibited a positive tendency, whereas forest edges and woodlands and hedgerows showed negative tendencies. No clear association was detected for winter cereals or other open-field habitats (Fig. 2).

**Figure 2.**
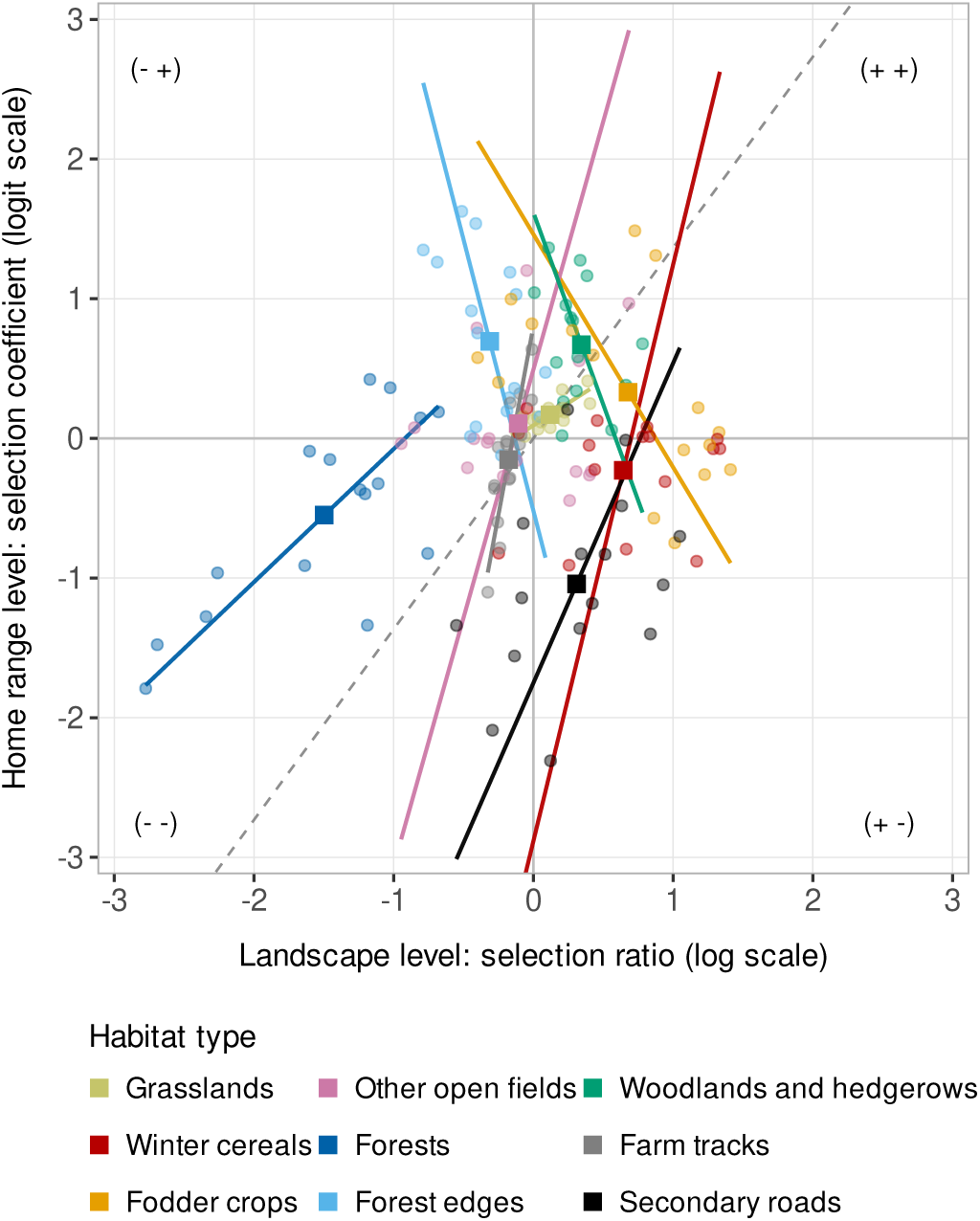
Association between landscape and home range habitat selection coefficients of released European hares (Lepus europaeus) across habitat types, from summer 2021 to spring 2023 in the Jura Mountains, France (52 hares, 108 seasonal home ranges). Selection coefficients at the landscape and home range levels were centred prior to analysis. Dashed lines show the overall association across habitat types, while coloured solid lines show habitat-specific relationships. All fitted lines are predictions of Deming regressions.

**Table 2.**
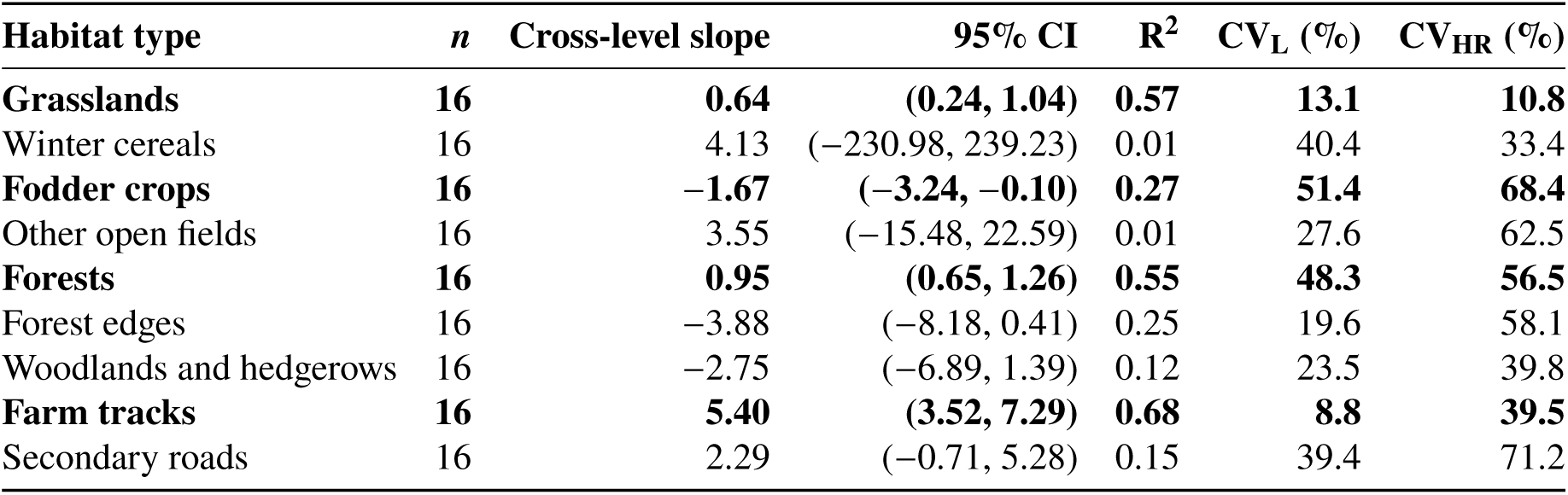
Cross-level coupling and temporal variability of landscape and home range habitat selection across habitat types in released European hares (*Lepus europaeus*), from summer 2021 to spring 2023 in the Jura Mountains, France (52 hares, 108 seasonal home ranges). The landscape-level selection was quantified as log-transformed selection ratios (log(SR)) and the home range-level selection as logit-scale selection coefficients (*β*); estimates were centred prior to analysis and 95% confidence intervals are reported. Slopes > 0 indicate concordant coupling between levels, whereas slopes < 0 indicate opposing selection across levels. *n* denotes the number of season-year-sex estimates used per habitat type. Deming regression accounted for uncertainty in both axes, with the error ratio estimated from the mean squared standard errors of estimates at the landscape and home range levels. CV_L_ and CV_HR_: coefficients of variation (*σ*/*μ*) of the predicted SRs (landscape) and ORs (home range) across the eight monitoring seasons and both sexes. Built-up areas (excluded from the home range analysis) had a landscape-level CV of 105%, owing to near-zero SRs. Bold rows indicate habitat categories for which the 95% confidence interval of the slope does not overlap zero.

### Site fidelity and relative temporal variability between orders

Within individuals, seasonal home range overlap was moderate using polygon overlap (SI = _0.52_ 0.56 _0.61_) but higher when accounting for intensity of space use (BC = _0.66_ 0.71 _0.76_). Consistent with our assumption that temporal variability would be more constrained at the landscape level than within home ranges, coefficients of variation of predicted selection across the eight seasons were higher at the home range level than at the landscape level for seven of the nine habitat types (Table 2). Only grasslands and winter cereals were slightly less variable within home ranges (Table 2).

The 48 transitions between consecutive seasonal home ranges (27 hares, 33 transitions within a land-cover map year) showed high fidelity (mean BC = 0.79; 10th–90th percentiles 0.65–0.92; range 0.16–0.97). Changes in landscape-level selection were only weakly compensated at the home range level (*c* = −0.06±0.14 within a map year; −0.13±0.13 for all transitions). Contrary to our prediction, the use of a habitat by European hares tracked its availability within the home range. Within a map year, the sign of the interaction between availability change and fidelity was in the direction predicted by H2 (*δ* = 0.77 ± 0.50 SE; parametric bootstrap: *p* = 0.068; two-stage weighted regression: *p* = 0.027). One single transition of low fidelity (BC = 0.16) however carried 24% of the weight in the two-stage analysis (Supporting information). Removing this transition reversed the sign of the interaction, which was no longer significant (*δ* = −1.14 ± 1.00 SE; bootstrap *p* = 0.18; two-stage *p* = 0.42). The absence of consistent interactive effects was further supported by the unweighted analyses and the analyses of all transitions (*δ* = 0.16 ± 0.69 SE; *p* ≥ 0.14). Using each individual’s CV of habitat availability as a habitat-specific measure of fidelity led to the same conclusion (interaction = −0.004 ± 0.37 SE, *p* = 0.99 within a map year). A complementary habitat-level analysis also showed no stronger cross-level coupling for habitat types whose availability varied more between seasons (Supporting information). Overall, H2 was not supported.

### Environmental drivers of temporal variation

At the landscape level, all models incorporating environmental covariates ranked below the best model according to BIC (Table 1a), although seasonal precipitation, NDVI, and temperature accounted for R^2^_ANODEV_ values of 0.24, 0.10, and 0.08, respectively, of the temporal deviance in habitat selection among habitat types. At the home range level, by contrast, the best model retained temporal environmental covariates, with habitat-specific responses (Table 1b). Seasonal temperature, seasonal precipitation, and NDVI accounted for R^2^_ANODEV_ values of 0.35, 0.19, and 0.12, respectively, of the temporal deviance in habitat selection across habitat types. The best-supported model, including additive interactions between habitat type and both temperature and precipitation, was strongly favoured over the second-best model and was retained for interpretation (Table 1b). We detected weather effects at the home range level only, differing between habitats (Fig. 3). Relative selection for forests declined sharply with increasing temperature, whereas selection for other open fields, farm tracks and secondary roads increased markedly (Fig. 3a). In contrast, no clear temperature effect was detected for grasslands, winter cereals, fodder crops, forest edges or woodlands and hedgerows (Fig. 3a). On the other hand, relative selection increased with precipitation in grasslands, forests, and forest edges, whereas it declined in other open fields, winter cereals, fodder crops, and secondary roads (Fig. 3b). Woodlands and hedgerows showed a moderate decrease with increasing precipitation, while no clear effect was detected for farm tracks (Fig. 3b; see Supporting information for slope estimates). H3 was therefore only partially supported because weather did not generate positive or negative association between levels.

**Figure 3.**
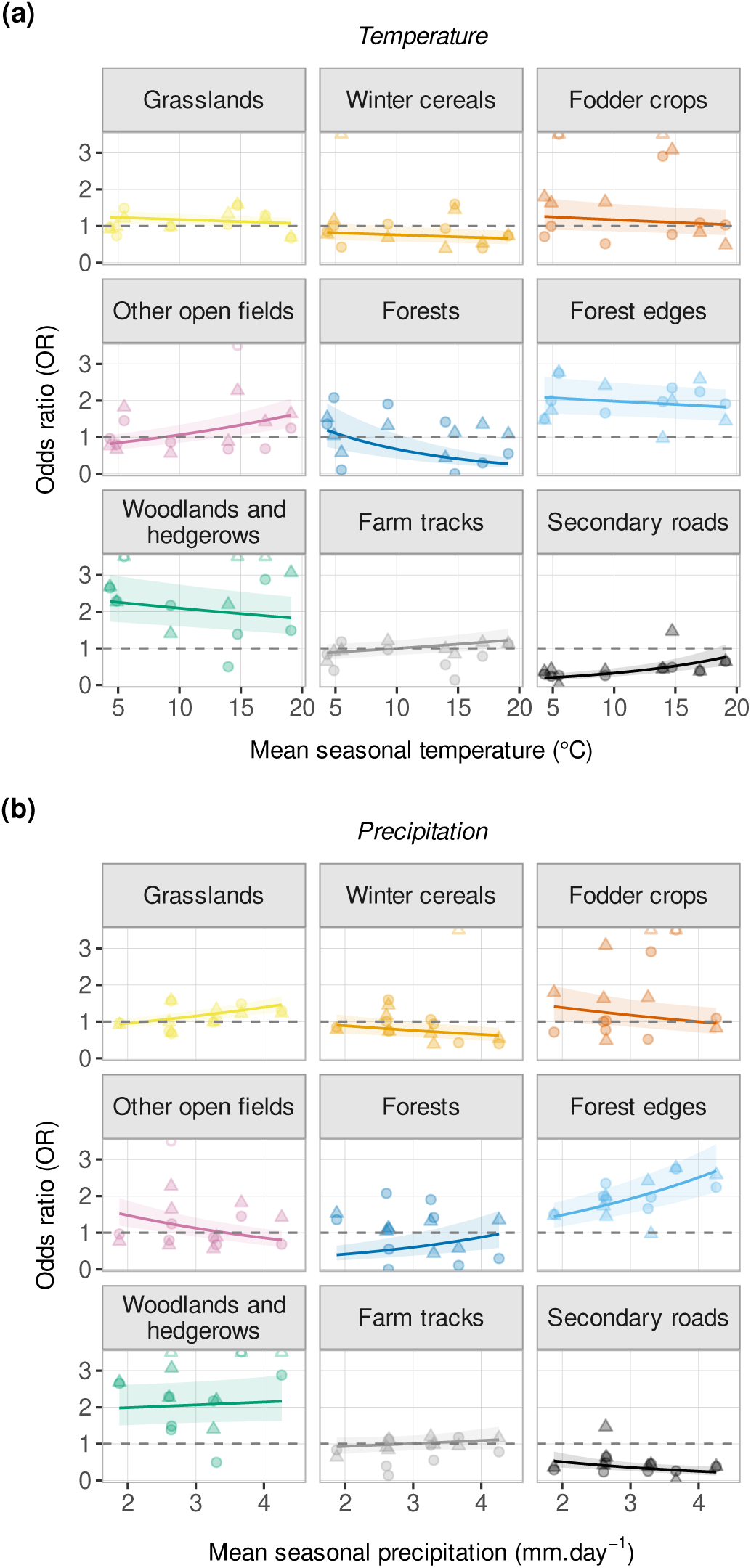
Home range habitat selection (odds ratios, ORs) of released European hare (*Lepus europaeus*) according to habitat types and (a) mean seasonal temperature and (b) mean seasonal precipitation, from summer 2021 to spring 2023 in the Jura Mountains, France (52 hares, 108 seasonal home ranges). Habitats with ORs > 1 are considered selected, whereas those with values < 1 are considered avoided. Lines and shaded ribbons show model-based ORs (estimated marginal means contrasts vs the grand mean) and the 95% confidence intervals. Triangles and circles represent empirical ORs computed for males and females, respectively.

## Discussion

The relationship between landscape- and home range-level selection was neither consistently reinforcing nor opposed, but habitat-dependent. Habitat selection by hares changed in opposite directions through time for fodder crops but in the same direction for grasslands, forests, and farm tracks. Three processes could account for these contrasting relationships: (*i*) hierarchical habitat availability, whereby limited or heterogeneous availability within the home range may promote stronger selection for particular patches (Van Moorter et al. 2016, Paolini et al. 2019), (*ii*) environmental conditions acting differently at each order (Dupke et al. 2017), and (*iii*) home range fidelity constraining second-order selection (Rettie and Messier 2000, Dupke et al. 2017). Although habitat availability may explain the negative relationship for fodder crops, the use of a habitat rather tracked its availability within the home range. Our results also suggest that environmental conditions acted differently at each order, and provide no support for a site-fidelity hypothesis.

Negative cross-level relationships may arise when a habitat more selected at the landscape level becomes more available within the home range, reducing the strength of subsequent fine-scale selection, whereas lower availability promotes stronger selection for the patches that remain accessible (Ofstad et al. 2019). Fodder crops fitted this mechanism best. Scarce in the landscape and selected at both levels, they showed highly variable selection over time (Table 2) and the only negative relationship relationship. Their patchy and seasonally variable availability may favour strong but changing selection at both levels, while greater incorporation during home range placement could reduce the strength of selection for the patches available within the home range (Reichlin et al. 2006, Santilli et al. 2024). Forest edges showed a similar negative tendency, although the mechanism is less clear. This habitat was avoided by males at the landscape level, possibly as a by-product of strong forest avoidance, but selected by females within home ranges, with a tendency in males.

Hares may not favour edge-rich areas when establishing home ranges but concentrated on particular edges once available, probably to seek shelter shelter and escape opportunities they provide (Petrovan et al. 2013, Naldi et al. 2020). Allowing sex differences to vary among seasons and habitats did not reverse the relationships for grasslands, farm tracks, forests and fodder crops, but confidence intervals widened and those for forests and fodder crops included zero (Appendix S2). Our results therefore hold, but our data had limited statistical power to detect these relationships at this resolution.

Positive cross-level relationships were, however, inconsistent with our initial expectation and may be observed when temporal changes in habitat value affect home range placement and within home range use in the same direction, outweighing the compensatory effect of habitat availability at the finer order. Grasslands, selected or used in proportion to availability, showed low temporal variability at both orders (Table 2) and parallel seasonal changes in selection. As they are mostly permanent, extensively grazed, and plant-diverse, grasslands likely provide stable year-round access to high-quality forage (Santilli et al. 2024), with variation in vegetation height also providing cover (Smith et al. 2004, Schai-Braun et al. 2015), so that small seasonal changes in their value would affect selection similarly at both orders. Forest selection was also congruent between levels. Forested habitats were avoided by hares at the landscape level (Hunter et al. 2026), and generally within home ranges, although somewhat less in autumn and winter. At this time of the year, forests may provide complementary shelter when cover in open habitats becomes limited (Tapper and Barnes 1986, Neumann et al. 2012). Such shelter value of forests could shift its selection at both orders in winter. The remaining habitats (*e.g.,* winter cereals, other open fields) showed little association between temporal changes at the two levels, (Table 2). This pattern could arise if selection at each level responds to different factors, such as seasonal resource quality and shelter availability (Rettie and Messier 2000, Dupke et al. 2017).

We hypothesised that home range fidelity would also shape cross-order relationships (Dupke et al. 2017, Kreling et al. 2021). By limiting shifts in home range placement, and hence temporal change in second-order selection, fidelity leaves third-order selection free to vary within an established home range even with little spatial reorganisation (Van Moorter et al. 2016, Kreling et al. 2021). At the extreme, perfect home range fidelity would leave little scope for temporal variation in second-order selection, and hence for covariation with third-order selection. Despite high seasonal overlap in home range extent and intensity of use, hares kept some flexibility in home range placement for second-order selection to vary across seasons, although less than third-order selection (Bennitt et al. 2014). Limited second-order variation did not, however, weaken cross-order relationships as predicted. For instance, selection for farm tracks varied little at the landscape level (Table 2) but was positively related to third-order selection. Contrary to our expectation, fidelity may constrain second-order variation without preventing cross-order relationships. The flexibility in site fidelity and habitat selection could allow seasonal trade-offs among habitat functions to be expressed at both orders, while habitat-specific requirements determine whether their temporal dynamics are antagonistic or concordant (Paolini et al. 2019).

Finally, environmental conditions offer another explanation to the varying relationships between selection orders we report. Weather variables did account for the temporal variation of habitat selection at the home range level but not at the landscape level. This observation is consistent with differences in behavioural responses to short-term environmental variability previously reported across hierarchical levels in other herbivores (Rettie and Messier 2000, Pape and Löffler 2015, Dupke et al. 2017). In the European hare, responses to temperature and precipitation differed among habitats, possibly reflecting their thermal, protective, and foraging functions (Milling et al. 2018, Mayer et al. 2019), through direct behavioural responses to weather (Gigliotti et al. 2018, Stiegler et al. 2023) or indirectly through changes in vegetation productivity, structure, and crop maturation (Rigoudy et al. 2024). Warmer conditions seemed to favour other open fields, farm tracks and secondary roads, while greater use of forests and forest edges under wetter conditions may indicate an increased value of protective cover under harsh environmental conditions (Oliveira-Santos et al. 2010, Hummel et al. 2017). Responses to precipitation also differed among open habitats, suggesting that weather effects depend on habitat function rather than following a simple open–wooded gradient.

## Conclusion

Because the released hares were captive-bred, the observed habitat selection pattern may differ from that of wild individuals because, for instance, of differences in familiarity with the landscape or anti-predator behaviour). Nevertheless, we investigated biological mechanisms by comparing temporal variation at the second- and third-order selection rather than absolute selection strength. Such relationships are unlikely to be strongly biased by animal origin since these cross-orders association should apply to any individuals. Taken together, our results show that cross-level relationships are a dynamic component of hierarchical habitat selection, and that selection observed at one spatial level cannot simply be extended to another (Rettie and Messier 2000, DeCesare et al. 2012, Dupke et al. 2017, Paolini et al. 2019). Future studies could quantify habitat composition and configuration within home ranges and integrate behavioural information (*e.g.,* foraging, resting, or sheltering) to determine whether cross-order relationships reflect the availability of complementary habitats, their functional use, or both. For a declining species such as the European hare (Smith et al. 2004, Gryz and Krauze-Gryz 2022), our findings support managing grassland-dominated landscapes (Zellweger-Fischer et al. 2011) as heterogeneous mosaics, with variation in vegetation structure, smaller fields, and diverse crop and semi-natural habitats (Smith et al. 2004, Mayer et al. 2018, Langhammer and Grimm 2020), that provide forage and cover across seasons (Tapper and Barnes 1986, Ullmann et al. 2023), rather than promoting individual habitat features in isolation.

## Supporting information

Supplementary figures, tables and additional analyses

## AI use statement

OpenAI ChatGPT was used by one of the authors to assist with R code development and debugging and as a discussion tool to examine statistical analyses and their possible interpretations. ChatGPT was also used to suggest formulations and assist with the linguistic refinement of individual sentences and passages throughout the manuscript. Google Labs Science (Literature Insights/Gemini for Science) was used by the same author to assist with literature searches and the identification of possible relevant scientific publications, and all publications considered for inclusion were checked against the original sources. No raw data or spatially explicit data from the study were provided to ChatGPT or Gemini. Anthropic Claude (Claude Code) was used by another author to generate Gitlab repository and verification of reported values and references. All AI-assisted code, analyses, and text were carefully reviewed and revised by the authors before inclusion in the manuscript. The authors retain full responsibility for the scientific content, interpretations, and conclusions.

## Acknowledgements

The authors thank Alexandre Tarral (GAEC Tarral, Aveyron, France) for breeding, rearing, and supplying the released hares. We are grateful to the FDC25 volunteers for their help with hare releases and field assistance. We thank Jean-Michel Bourbon (local hunter) and the XERIUS team, notably Virginie Delepoulle, for their support during fieldwork. The French National Geographic Institute (IGN) provided all geographical data. We are also deeply grateful to Bram van Moorter for helping us to understand the cross-level mechanisms in habitat selection theories.

## Funding

This research was funded by the OFB (French Biodiversity Office), FNC (National Federation of Hunters), FDC25 (Federation of Hunters of the Doubs Department), and Faune INNOV’ R&D.

## Conflict of interest disclosure

The authors declare that they comply with the PCI rule of having no financial conflicts of interest in relation to the content of the article. The authors declare non-financial conflicts of interest: T. Powolny and R. Renaude are working at the Fédération départementale des chasseurs du Doubs, which co-funded the study and ran the hare reinforcement programme; L. Berger, C. P. Henriot and M. Sage are employed by Faune INNOV’ R&D, a private wildlife research consulting company that co-funded the study and received funding from hunters’ federations.

## Ethics approval

This study complies with European and French national regulations on the protection of animals used for scientific purposes (Directive 2010/63/EU). Hares were fitted with external GPS collars without anaesthesia or surgery; under French regulations (note de service DGAL/SDSPA/N2013-8095), this procedure falls outside the scope of these regulations, so no ethics committee approval or project authorisation was required. The released hares were bred at a game farm (official number 12412) declared to the departmental agriculture and veterinary services of Aveyron, which adheres to the charter of the French game breeders’ union and undergoes an annual health inspection by an accredited veterinarian. The release of European hares is not subject to prefectural authorisation under article L424-11 of the French Environment Code.

## Data, scripts, code, and supplementary information availability

De-identified data and all code needed to reproduce the analyses are available at https://gitlab.in2p3.fr/christophe.bonenfant/hare-habitat-selecti on-code (version v1.0.3) and will be deposited in a versioned open repository with a DOI upon recommendation. We withheld raw GPS locations because disclosing spatially explicit information on hare locations could increase poaching risk. Raw data will be made available from the corresponding author on reasonable request. Supplementary information is provided as a separate file.

## Author contributions

MS, RR, and LB designed the fieldwork. MS, TP, CPH, RR, and CB supervised the project. LB, MS, and RR performed the experiments. LB, CPH, and CB analysed the data. LB, CPH, and CB wrote the manuscript with the support of TP and MS.

## Notes

https://gitlab.in2p3.fr/christophe.bonenfant/hare-habitat-selection-code

### Competing Interest Statement

The authors declare non-financial conflicts of interest: T. Powolny and R. Renaude are working at the Federation departementale des chasseurs du Doubs, which co-funded the study and ran the hare reinforcement programme; L. Berger, C.P. Henriot and
M. Sage are employed by Faune INNOV' R\&D, a private wildlife research consulting company that co-funded the study and received funding from hunters' federations.

