## Supplementary figures, tables and additional analyses for "Cross-level correlation and temporal dynamics of hierarchical habitat selection"

### Table of Contents:

| | |
|---|---|
| **Figure S1. Sensitivity of home range habitat selection estimates to RSF formulation following the Muff et al. (2020) framework.** | Page 3 |
| **Figure S2. 95% wAKDE seasonal home ranges of the 52 European hares released in 2021 and 2022 in the Jura Mountains, France, used in the habitat selection analyses.** | Page 4 |
| **Figure S3A. Correlation matrix of predicted selection ratios (habitat selection at the landscape level) among habitat types in the European hare.** | Page 5 |
| **Figure S3B. Correlation matrix of predicted odds ratios (habitat selection at the home range level) among habitat types in the European hare.** | Page 6 |
| **Figure S4. Cross-level coupling of habitat selection as a function of the seasonal variability of habitat availability within home ranges.** | Page 7 |
| **Figure S5. Cross-level coupling of habitat selection for each transition between consecutive seasonal home ranges as a function of fidelity.** | Page 8 |
| **Table S1. Definition and composition of habitat types used in the habitat selection analysis for the European hare.** | Page 9 |
| **Table S2. Comparison of random-effects structures in home range-level habitat selection models.** | Page 10 |
| **Table S3. Sensitivity analysis of cross-scale associations using ordinary linear models.** | Page 11 |
| **Table S4A. Coefficients of the best landscape-level habitat selection model.** | Page 12 |
| **Table S4B. Coefficients of the best home range-level habitat selection model.** | Page 13 |
| **Table S5. Habitat-specific effects of temperature and precipitation on home range habitat selection.** | Page 14 |
| **Table S6. Seasonal variability of habitat availability within home ranges and cross-level coupling of habitat selection, by habitat type.** | Page 15 |
| **Table S7. Sensitivity of the cross-level slopes to the error model and to sex differences among seasons.** | Page 17 |
| **Appendix S1. Habitat-level analysis of site fidelity and cross-level coupling (H2).** | Page 16 |
| **Appendix S2. Sensitivity of the cross-level slopes to sex differences among seasons.** | Page 17 |

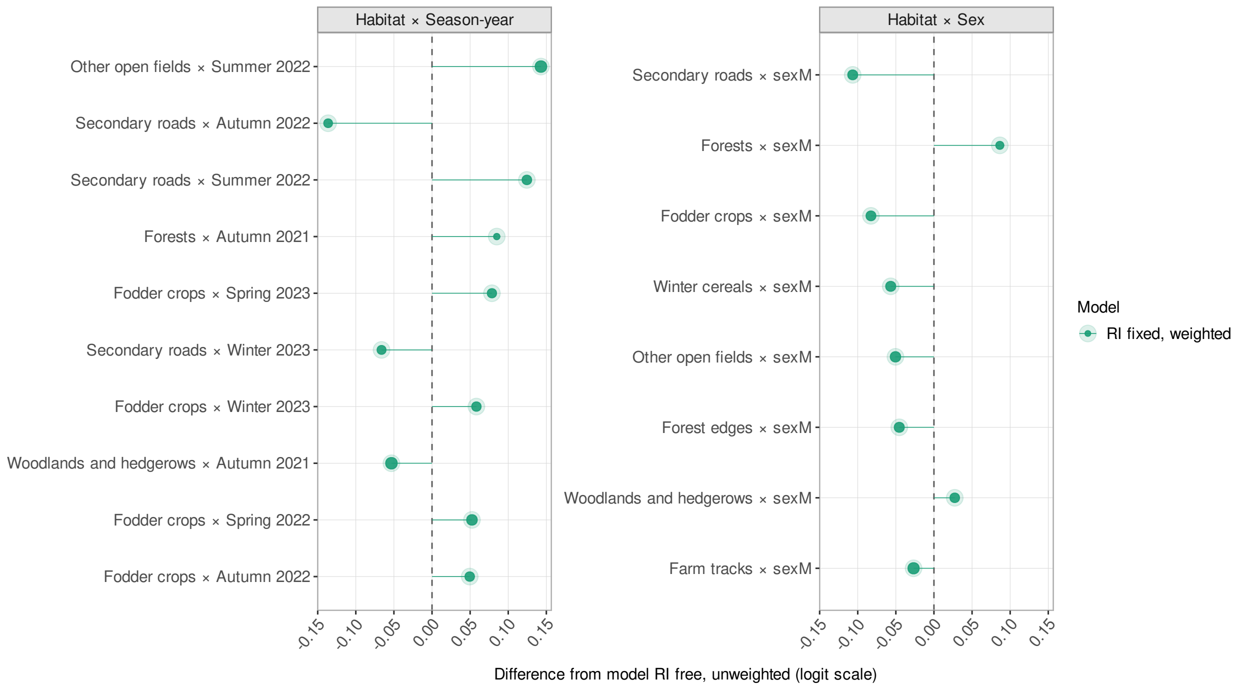


**Figure S1.** Sensitivity of home range habitat selection estimates to RSF formulation following the Muff et al. (2020) framework. RI refers to the treatment of individual-specific random intercepts (tag_id), which were either freely estimated or fixed to a large value (SD = 1000) to approximate a conditional likelihood and avoid shrinkage, following the framework of Muff et al. (2020). In addition, weights were set to 1 for used locations and 1000 for available locations to approximate an inhomogeneous Poisson point process (IPP). All models were fitted with an identical random-effects structure, including an individual-specific intercept and uncorrelated habitat-specific random slopes `[(1 | tag_id) + (0 + habitat || tag_id)]`, ensuring that differences among models arise solely from the treatment of intercept variance and weighting. Coefficient differences are expressed relative to the unweighted model with freely estimated random intercept variance. Point size is inversely proportional to the width of the 95% confidence interval, with larger points indicating more precise estimates.

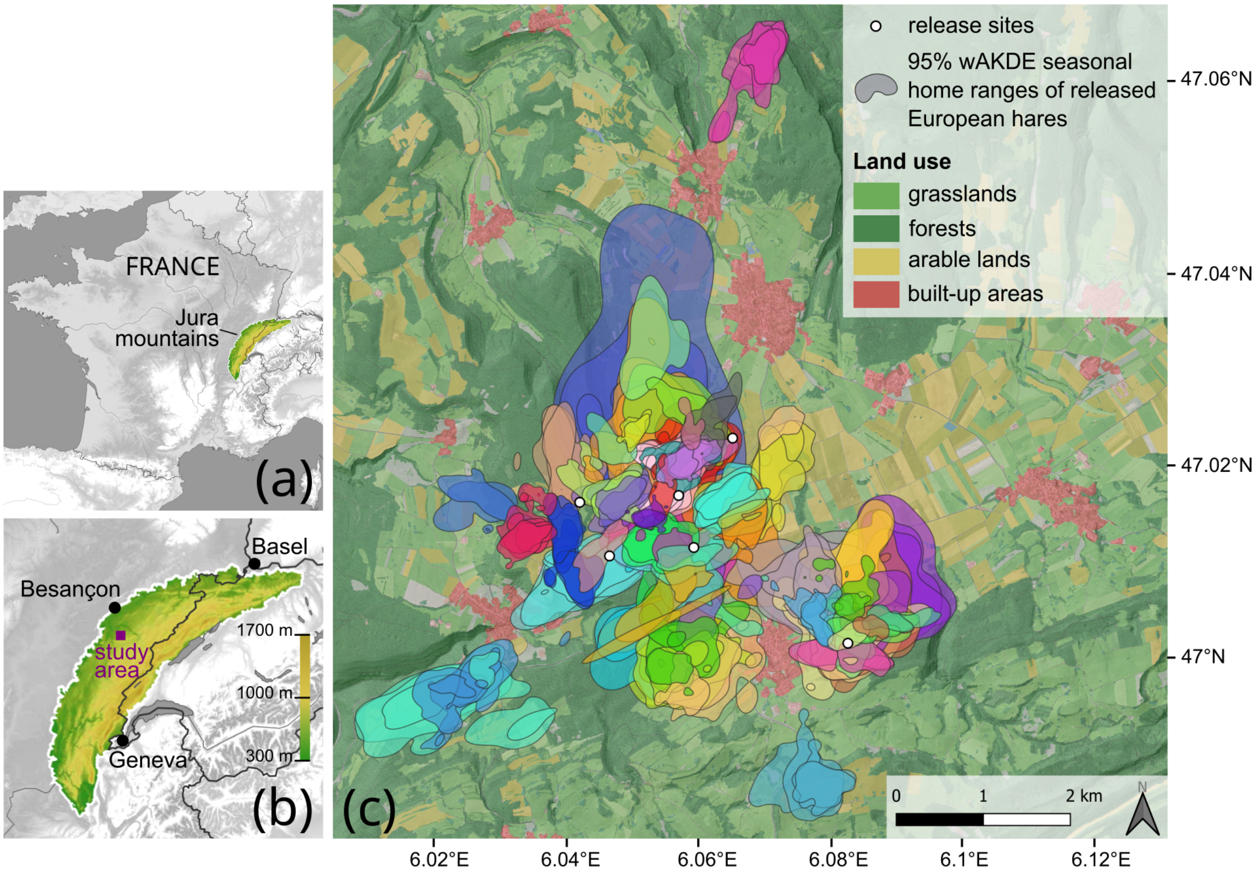


**Figure S2.** 95% wAKDE (weighted autocorrelated kernel density estimation) home ranges (HRs) of the 52 European hares released in 2021 and 2022 in the Jura Mountains, France, used in the habitat selection analyses. (a) Location of the Jura Mountains in France; (b) location of the study area in the Jura Mountains, with elevation; (c) home ranges and release sites over land use. The six sites where the 103 European hares were released in early July 2021 and early September 2022 are located by white circles on the map. The background is a combination of land use and relief. The coordinate reference system is WGS84.

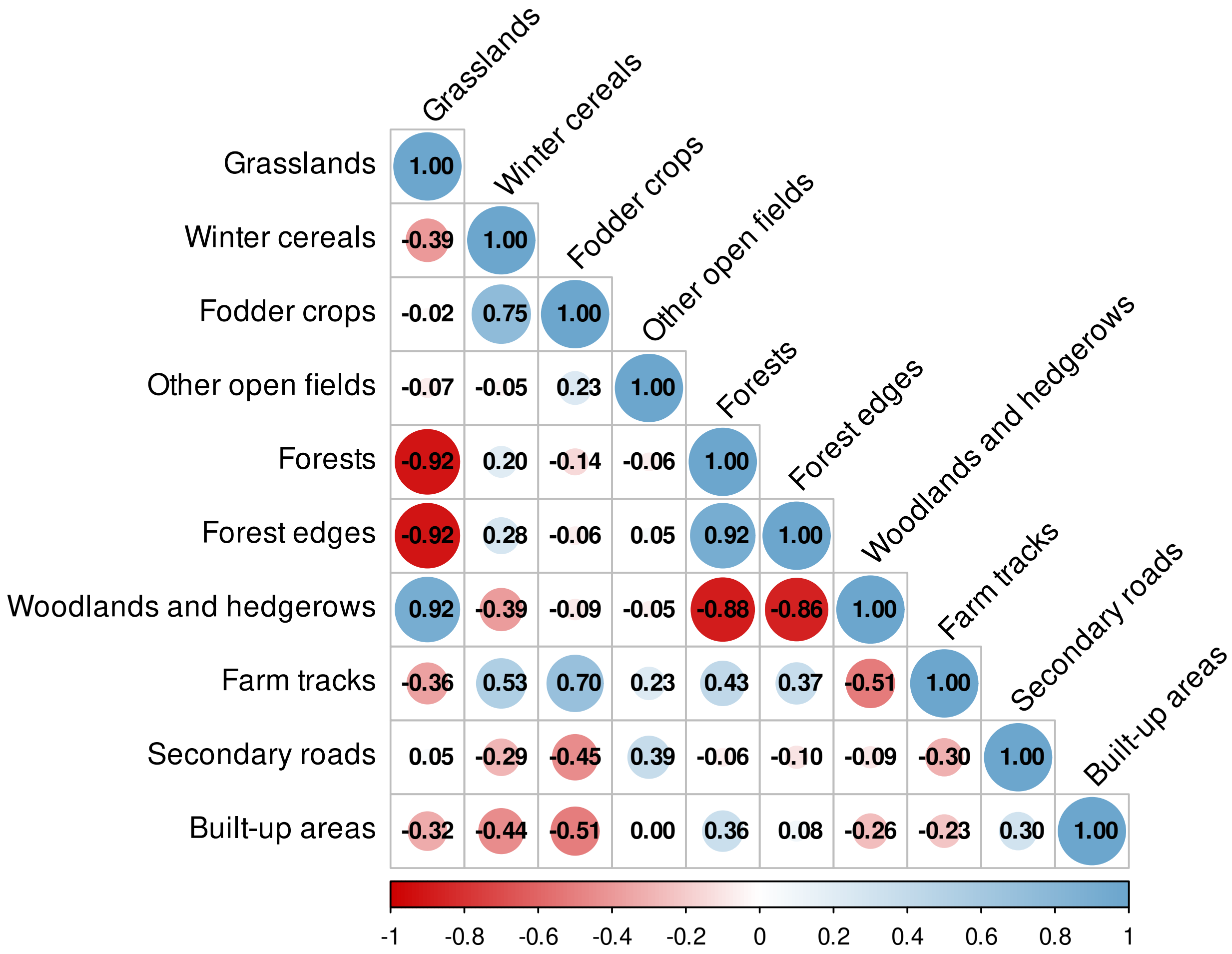


**Figure S3A.** Correlation matrix of predicted selection ratios (habitat selection at the landscape level) for the different habitat types in the European hare (*Lepus europaeus*). Each cell represents the Pearson correlation coefficient between predicted selection ratios for two habitat types across season-years. Predictions from a model fitted to 52 hares (108 seasonal home ranges).

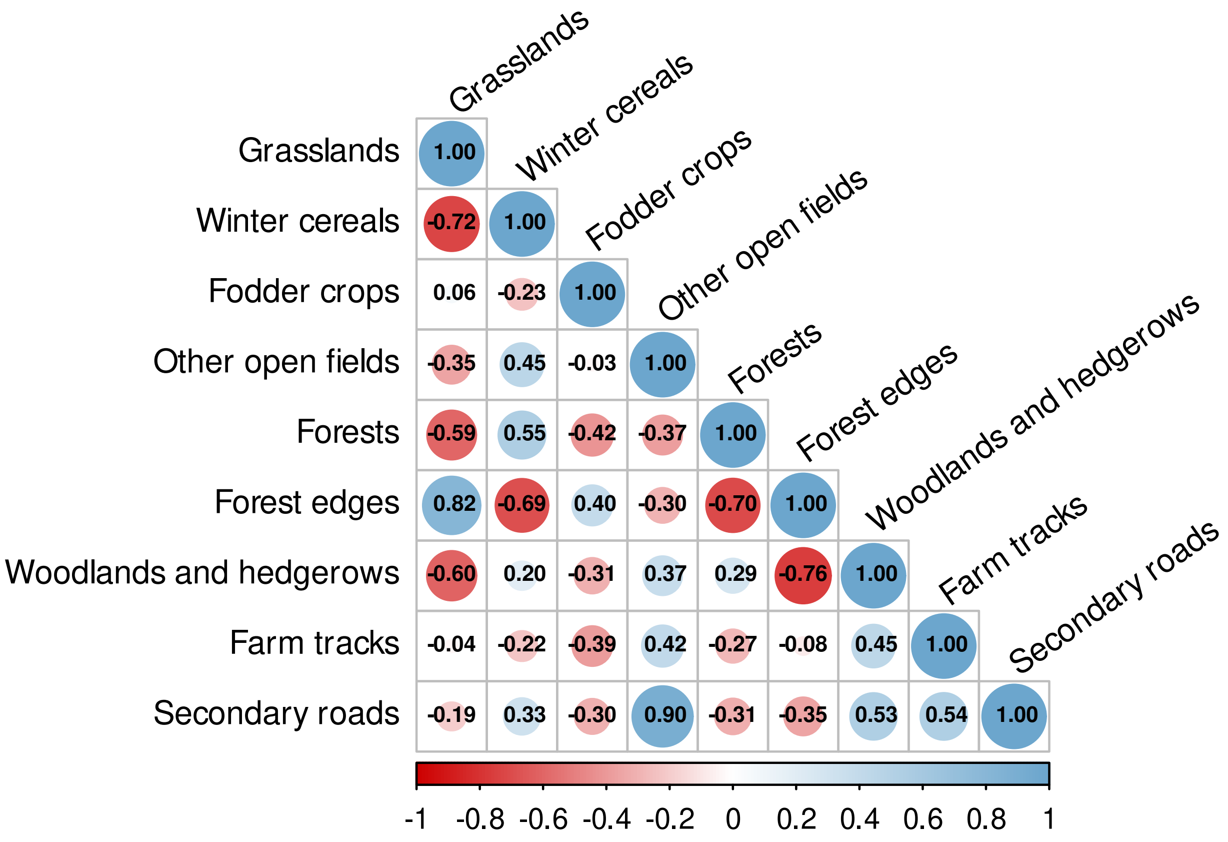


**Figure S3B.** Correlation matrix of predicted odds ratios (habitat selection at the home range level) among habitat types in the European hare (*Lepus europaeus*). Each cell represents the Pearson correlation coefficient between predicted odds ratios for two habitat types across season-year combinations. Predictions from a model fitted to 52 hares (108 seasonal home ranges).

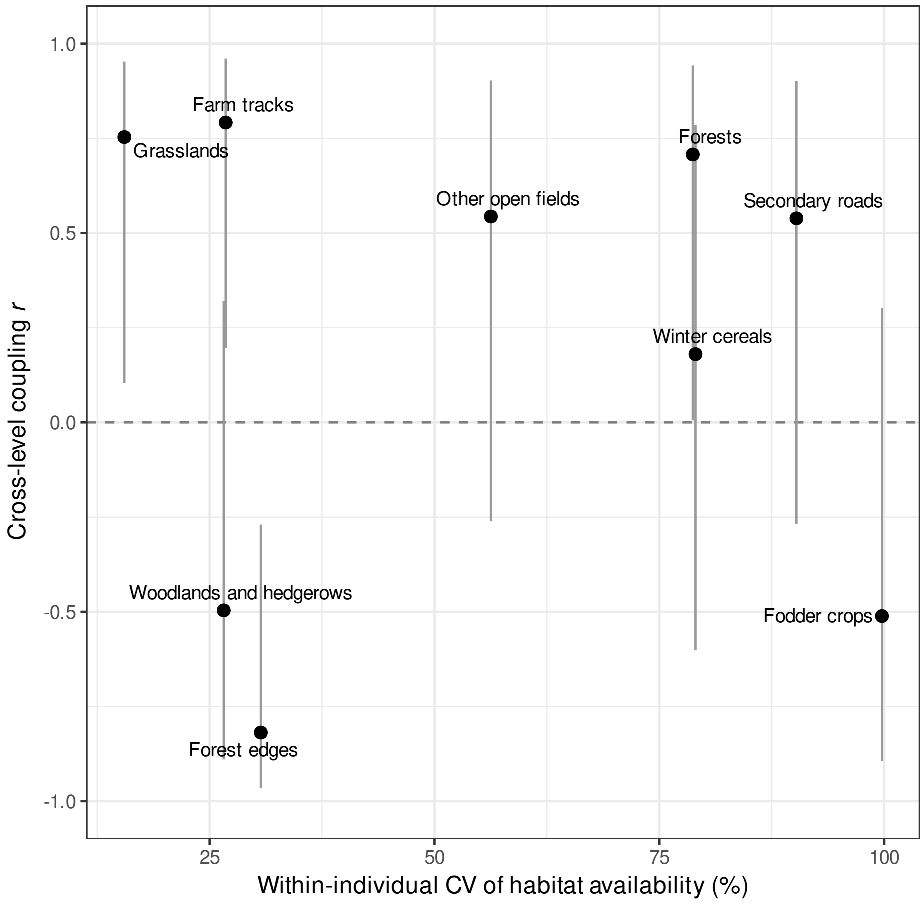


**Figure S4.** Cross-level coupling of habitat selection as a function of the seasonal variability of habitat availability within home ranges, by habitat type, for European hares (*Lepus europaeus*) released in the Jura Mountains, France, from summer 2021 to spring 2023 (52 hares). Coupling is the Pearson correlation (*r*) between landscape- and home range-level seasonal predictions of selection, centred within sex (eight season-years); bars show 95% confidence intervals from Fisher's *z* transformation. Variability is the within-individual coefficient of variation (CV) of the proportion of each habitat type within seasonal home ranges, averaged over the 30 individuals monitored for at least two seasons (Table S6, Appendix S1).

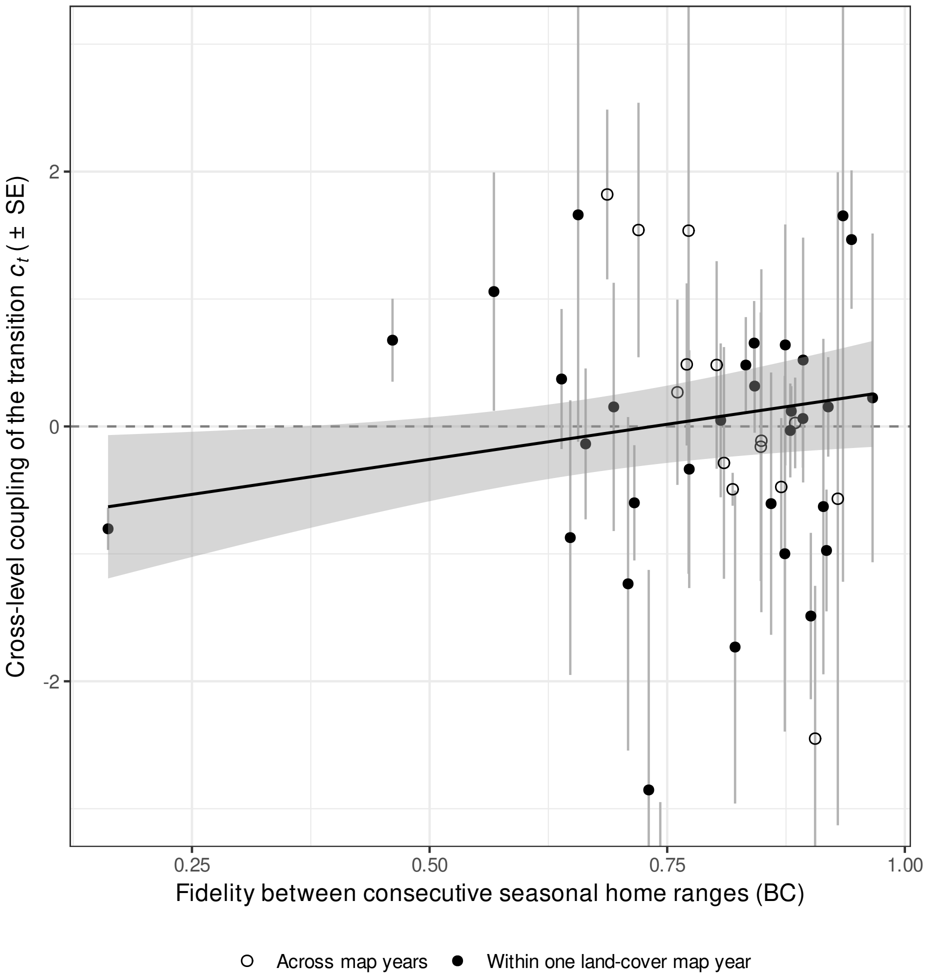


**Figure S5.** Cross-level coupling of habitat selection ($c_t \pm$ SE) for each transition between two consecutive seasonal home ranges, as a function of fidelity (Bhattacharyya coefficient, BC, between the two utilisation distributions), for European hares (*Lepus europaeus*) released in the Jura Mountains, France, from summer 2021 to spring 2023 (48 transitions from 27 hares). $c_t = 0$: use changes in proportion to availability (no coupling); $c_t = -1$: full compensation within the home range. Filled symbols: transitions within one land-cover map year ($n = 33$, main analysis); open symbols: transitions across map years. Line and shaded band: regression of $c_t$ on fidelity weighted by the inverse sampling variance of $c_t$, with its 95% confidence interval, for transitions within one map year. The leftmost transition (BC = 0.16) carries 24% of the weight of this regression; without it the relationship reverses sign and is not significant. Three transitions with $|c_t| > 3$ fall outside the plotted range.

**Table S1.** Definition and composition of habitat types used in the habitat selection analysis for the European hare (*Lepus europaeus*), resulting from the aggregation of original land-cover classes.

| **Habitat type** | **Dominant land-cover classes** |
|---|---|
| *Open habitats* | |
| Grasslands | – Permanent meadow (> 6 years)<br>– Long-term rotational meadow (> 6 years)<br>– Temporary meadow (< 5 years) |
| Winter cereals | – Winter barley<br>– Winter wheat<br>– Winter mixed cereals |
| Fodder crops | – Legume-forage grass mixture ($\leq$ 5 years)<br>– Alfalfa |
| Other open fields | – Undetermined open fields / field margins<br>– Maize<br>– Winter rapeseed |
| *Tree/shrub dominated habitats* | |
| Forests | – Closed broadleaf forest (relative broadleaf cover > 75%)<br>– Closed coniferous forest (relative coniferous cover > 75%)<br>– Closed mixed forest (25–75% relative broadleaf cover) (area > 0.5 ha, absolute tree cover > 40%)<br>– Open forest (area > 0.5 ha, 10–40% absolute tree cover) |
| Forest edges | – Forest-open habitat ecotones (50 m inside forest) |
| Woodlands and hedgerows | – Woodland (area < 0.5 ha)<br>– Hedgerow (width < 20 m, length > 25 m) |
| *Anthropogenic features* | |
| Farm tracks | – Agricultural tracks and dirt roads (3 m buffer) |
| Secondary roads | – Secondary road network (6 m buffer) |
| Built-up areas | – Residential and commercial areas (30 m buffer)<br>– Farmyards (no buffer) |

**Table S2.** Comparison of random-effects structures in home range-level habitat selection models. Both models include time since release and habitat type as fixed effects. Models were ranked using the Bayesian information criterion (BIC), with lower values indicating better fit. $\Delta$BIC is calculated relative to the best-supported model. No evidence of singularity was detected for either model.

| Model | df | BIC | $\Delta$BIC | Singular fit |
|---|---|---|---|---|
| Random intercept + random slopes<br>`(1 \| tag_id) + (0 + habitat \|\| tag_id)` | 20 | 325 941 | 0 | No |
| Random intercept only<br>`(1 \| tag_id)` | 11 | 333 206 | 7264 | No |

**Table S3.** Sensitivity analysis of cross-scale associations using ordinary linear models. One linear model was fitted independently for each habitat type using centred raw coefficients derived from the GLMM outputs. Reported values correspond to slope estimates, 95% confidence intervals (CI), $p$-values, and coefficients of determination ($R^2$). Bold rows indicate slopes significantly different from 0. Conclusions matched those of the Deming regressions (Table 2 of the main text) for all habitat types except forest edges, whose slope was marginally significant here but not under Deming regression (95% CI: $-8.18$, 0.41).

| Habitat type | Slope | 95% lower CI | 95% upper CI | *p*-value | $R^2$ |
|---|---|---|---|---|---|
| **Grasslands** | **0.51** | **0.26** | **0.77** | **<0.01** | **0.57** |
| Winter cereals | 0.07 | −0.36 | 0.51 | 0.72 | 0.01 |
| **Fodder crops** | **−0.55** | **−1.07** | **−0.03** | **0.04** | **0.27** |
| Other open fields | 0.13 | −0.47 | 0.72 | 0.66 | 0.01 |
| **Forests** | **0.76** | **0.37** | **1.16** | **<0.01** | **0.55** |
| **Forest edges** | **−1.20** | **−2.39** | **−0.01** | **0.048** | **0.25** |
| Woodlands and hedgerows | −0.67 | −1.71 | 0.37 | 0.19 | 0.12 |
| **Farm tracks** | **3.82** | **2.31** | **5.32** | **<0.01** | **0.68** |
| Secondary roads | 0.57 | −0.20 | 1.34 | 0.14 | 0.15 |

**Table S4A.** Coefficients of the best landscape-level habitat selection model. CI: confidence interval. Model fitted to 52 hares (108 seasonal home ranges).

| Term | Estimate | 95% lower CI | 95% upper CI |
|---|---|---|---|
| Intercept | 0.127 | 0.036 | 0.219 |
| *Main effects* | | | |
| Time since release | 0.010 | −0.030 | 0.049 |
| Sex (Male) | 0.182 | 0.061 | 0.304 |
| Habitat (Winter cereals) | 1.019 | 0.641 | 1.396 |
| Habitat (Fodder crops) | 1.035 | 0.529 | 1.542 |
| Habitat (Other open fields) | 0.266 | 0.021 | 0.512 |
| Habitat (Forests) | −1.482 | −1.855 | −1.108 |
| Habitat (Forest edges) | −0.154 | −0.331 | 0.023 |
| Habitat (Woodlands and hedgerows) | 0.265 | 0.089 | 0.441 |
| Habitat (Farm tracks) | −0.222 | −0.375 | −0.068 |
| Habitat (Secondary roads) | 0.536 | 0.155 | 0.916 |
| Habitat (Built-up areas) | −1.928 | −2.768 | −1.087 |
| *Interaction: habitat × sex* | | | |
| Winter cereals × Male | −0.719 | −1.254 | −0.185 |
| Fodder crops × Male | −0.604 | −1.318 | 0.109 |
| Other open fields × Male | −0.891 | −1.245 | −0.536 |
| Forests × Male | 0.202 | −0.285 | 0.688 |
| Forest edges × Male | −0.452 | −0.694 | −0.210 |
| Woodlands and hedgerows × Male | −0.072 | −0.309 | 0.165 |
| Farm tracks × Male | −0.103 | −0.310 | 0.105 |
| Secondary roads × Male | −0.529 | −1.063 | 0.005 |
| Built-up areas × Male | 0.304 | −0.792 | 1.400 |

**Table S4B.** Coefficients of the best home range-level habitat selection model. CI: confidence interval. Model fitted to 52 hares (108 seasonal home ranges).

| **Term** | **Estimate** | **95% lower CI** | **95% upper CI** |
|---|---|---|---|
| Intercept | −2.624 | −2.746 | −2.502 |
| *Main effects* | | | |
| Sex (Male) | 0.083 | −0.087 | 0.253 |
| Precipitation | 0.039 | 0.018 | 0.060 |
| Temperature | 0.017 | −0.004 | 0.038 |
| Time since release | 0.013 | −0.002 | 0.028 |
| Habitat (Winter cereals) | −0.450 | −0.859 | −0.041 |
| Habitat (Fodder crops) | −0.075 | −0.574 | 0.424 |
| Habitat (Other open fields) | −0.119 | −0.463 | 0.225 |
| Habitat (Forests) | −0.815 | −1.602 | −0.027 |
| Habitat (Forest edges) | 0.653 | 0.306 | 0.999 |
| Habitat (Woodlands and hedgerows) | 0.470 | 0.051 | 0.889 |
| Habitat (Farm tracks) | −0.214 | −0.512 | 0.083 |
| Habitat (Secondary roads) | −1.291 | −1.807 | −0.775 |
| *Interaction: habitat × sex* | | | |
| Winter cereals × Male | 0.013 | −0.545 | 0.571 |
| Fodder crops × Male | 0.136 | −0.543 | 0.814 |
| Other open fields × Male | 0.121 | −0.360 | 0.601 |
| Forests × Male | 0.366 | −0.680 | 1.412 |
| Forest edges × Male | −0.261 | −0.753 | 0.230 |
| Woodlands and hedgerows × Male | 0.206 | −0.377 | 0.790 |
| Farm tracks × Male | 0.144 | −0.271 | 0.559 |
| Secondary roads × Male | 0.159 | −0.508 | 0.826 |
| *Interaction: habitat × temperature* | | | |
| Winter cereals × Temperature | −0.029 | −0.078 | 0.019 |
| Fodder crops × Temperature | −0.018 | −0.108 | 0.072 |
| Other open fields × Temperature | 0.315 | 0.231 | 0.400 |
| Forests × Temperature | −0.517 | −0.583 | −0.451 |
| Forest edges × Temperature | 0.003 | −0.032 | 0.038 |
| Woodlands and hedgerows × Temperature | −0.029 | −0.079 | 0.020 |
| Farm tracks × Temperature | 0.183 | 0.082 | 0.285 |
| Secondary roads × Temperature | 0.593 | 0.366 | 0.821 |
| *Interaction: habitat × precipitation* | | | |
| Winter cereals × Precipitation | −0.253 | −0.312 | −0.194 |
| Fodder crops × Precipitation | −0.257 | −0.361 | −0.154 |
| Other open fields × Precipitation | −0.336 | −0.418 | −0.253 |
| Forests × Precipitation | 0.135 | 0.053 | 0.218 |
| Forest edges × Precipitation | 0.053 | 0.017 | 0.090 |
| Woodlands and hedgerows × Precipitation | −0.112 | −0.160 | −0.063 |
| Farm tracks × Precipitation | −0.080 | −0.193 | 0.033 |
| Secondary roads × Precipitation | −0.398 | −0.615 | −0.182 |

**Table S5.** Habitat-specific effects of temperature and precipitation on home range habitat selection. Reported values correspond to model-derived slope estimates ($\beta$) and their 95% confidence intervals (CI). Temperature effects represent the change in habitat selection associated with a +5 °C increase in mean seasonal temperature, whereas precipitation effects represent the change associated with a +0.72 mm day$^{-1}$ increase in mean seasonal precipitation. Estimates were calculated at mean values of the other covariates.

| **Habitat type** | **Trend ($\beta$)** | **95% lower CI** | **95% upper CI** |
|---|---|---|---|
| *Mean seasonal temperature* | | | |
| Grasslands | 0.01 | 0.00 | 0.03 |
| Winter cereals | −0.01 | −0.05 | 0.03 |
| Fodder crops | 0.00 | −0.08 | 0.08 |
| Other open fields | 0.29 | 0.22 | 0.36 |
| Forests | −0.44 | −0.49 | −0.38 |
| Forest edges | 0.02 | −0.01 | 0.04 |
| Woodlands and hedgerows | −0.01 | −0.05 | 0.03 |
| Farm tracks | 0.17 | 0.09 | 0.26 |
| Secondary roads | 0.53 | 0.33 | 0.73 |
| *Mean seasonal precipitation* | | | |
| Grasslands | 0.04 | 0.02 | 0.06 |
| Winter cereals | −0.21 | −0.27 | −0.16 |
| Fodder crops | −0.22 | −0.32 | −0.12 |
| Other open fields | −0.29 | −0.37 | −0.21 |
| Forests | 0.17 | 0.09 | 0.25 |
| Forest edges | 0.09 | 0.06 | 0.12 |
| Woodlands and hedgerows | −0.07 | −0.12 | −0.03 |
| Farm tracks | −0.04 | −0.15 | 0.07 |
| Secondary roads | −0.35 | −0.57 | −0.14 |

**Table S6.** Seasonal variability of habitat availability within home ranges and cross-level coupling of habitat selection, by habitat type, for European hares (*Lepus europaeus*) released in the Jura Mountains, France, from summer 2021 to spring 2023 (52 hares, 108 seasonal home ranges). Availability: proportion of random available points falling in each habitat type within each seasonal home range (mean over home ranges). $CV_{within}$: within-individual coefficient of variation of availability across seasons, averaged over the $n$ individuals monitored for at least two seasons in which the habitat type occurred (30 individuals in total). $CV_{pop}$: coefficient of variation of the population mean availability across the eight season-years. *r*: Pearson correlation between landscape- and home range-level seasonal predictions of selection, centred within sex (eight season-years), with its 95% confidence interval (CI) from Fisher's *z* transformation.

| Habitat type | Availability (%) | $CV_{within}$ (%) | *n* | $CV_{pop}$ (%) | *r* | 95% CI |
|---|---|---|---|---|---|---|
| Grasslands | 52.3 | 15.5 | 30 | 13.6 | 0.75 | (0.10, 0.95) |
| Winter cereals | 11.6 | 79.0 | 26 | 41.4 | 0.18 | (−0.60, 0.79) |
| Fodder crops | 3.4 | 99.7 | 21 | 52.6 | −0.51 | (−0.89, 0.30) |
| Other open fields | 4.9 | 56.3 | 30 | 22.1 | 0.54 | (−0.26, 0.90) |
| Forests | 6.2 | 78.7 | 28 | 49.5 | 0.71 | (0.01, 0.94) |
| Forest edges | 13.7 | 30.7 | 30 | 19.7 | −0.82 | (−0.97, −0.27) |
| Woodlands and hedgerows | 5.0 | 26.6 | 30 | 23.0 | −0.50 | (−0.89, 0.32) |
| Farm tracks | 1.8 | 26.8 | 30 | 7.6 | 0.79 | (0.20, 0.96) |
| Secondary roads | 1.1 | 90.2 | 27 | 40.2 | 0.54 | (−0.27, 0.90) |

### Appendix S1. Habitat-level analysis of site fidelity and cross-level coupling (H2)

**Methods.** We also examined the effect of site fidelity at the habitat level, testing whether cross-level coupling was weaker for habitat types whose availability within home ranges was more stable between seasons. We measured the composition of each seasonal home range as the proportion of available points in each habitat type and computed two metrics to measure its seasonal variability: the within-individual coefficient of variation (CV$_i$), averaged over the individuals monitored for at least two seasons ($n = 30$), and the CV of the population mean across the eight season-years. Because the Deming slope cannot be compared among habitat types, we measured coupling with the Pearson correlation ($r$) between landscape- and home range-level seasonal marginal means of each habitat type. We centred the marginal means within sex before computing $r$, so that differences in habitat selection between sexes did not contribute to the temporal coupling of interest; as neither model included a sex $\times$ season-year interaction, centred values were identical for both sexes and $r$ rested on eight season-years (95% confidence intervals from Fisher's $z$ transformation). We related $r$ and $|r|$ to both CVs with Spearman rank correlations ($n = 9$ habitat types).

**Results.** Habitat availability within home ranges varied little between seasons for grasslands (within-individual CV = 16%), woodlands and hedgerows (27%), farm tracks (27%) and forest edges (31%), but strongly for winter cereals (79%), forests (79%), secondary roads (90%) and fodder crops (100%) (Table S6). Cross-level coupling, measured by the correlation between seasonal landscape- and home range-level selection, was positive for grasslands ($r$ = ${}_{0.10}\,0.75\,{}_{0.95}$), forests (${}_{0.01}\,0.71\,{}_{0.94}$) and farm tracks (${}_{0.20}\,0.79\,{}_{0.96}$), negative for forest edges (${}_{-0.97}\,-0.82\,{}_{-0.27}$), and unclear for the other habitat types. Contrary to H2, coupling did not strengthen with the seasonal variability of habitat availability (Spearman $\rho = -0.37$, $p = 0.34$ for $r$; $\rho = -0.45$, $p = 0.23$ for $|r|$). If anything, the strongest associations, of either sign, occurred in the habitats whose availability was the most stable ($\rho = -0.67$, $p = 0.06$ between $|r|$ and the CV of the population mean; Fig. S4).

### Appendix S2. Sensitivity of the cross-level slopes to sex differences among seasons

**Methods.** The temporal models used for the cross-level analysis had no sex-by-season-year term. The difference between sexes was therefore constant across seasons, and the 16 estimates per habitat type were not independent, carrying the information of eight season-years. To evaluate this assumption, we refitted both models with a three-way interaction between habitat type, sex and season-year, all other terms unchanged. We excluded built-up areas at the landscape level, because they had no use in some season-years and are not part of the cross-level analysis. We compared the additive and three-way models fitted to the same data with likelihood ratio tests and BIC. We then fitted Deming regressions using the standard error of each estimate, with jackknife confidence intervals (*deming* R package; Therneau 2024), for each habitat type and pooling all habitat types. We applied these regressions to the additive models too, to separate the effect of the error model from that of the interaction.

**Results.** Allowing sex differences to vary among seasons and habitats did not improve the landscape-level model ($\chi^2 = 59.6$, $df = 63$, $p = 0.60$; ΔBIC = 373.8 in favour of the additive model). It improved the home range-level model ($\chi^2 = 361.7$, $df = 63$, $p < 0.001$), although BIC still favoured the additive model (ΔBIC = 471.4). Using the standard error of each estimate in the additive models did not change the conclusions of Table 2 of the main text (Table S7). In the three-way models, each estimate rested on 3 to 12 hares and most confidence intervals widened. Slopes remained positive for grasslands and farm tracks, with confidence intervals excluding zero. Slopes for forests and fodder crops kept their direction, but their confidence intervals included zero. Relationships for the other habitat types remained unclear, and the slope for secondary roads was unreliable because one estimate (autumn 2021, males) had a very large standard error. These results are consistent with a loss of precision rather than a change in the direction of the relationships, and indicate that our data had limited statistical power to estimate cross-level relationships when sex differences varied among seasons and habitats.

**Table S7.** Sensitivity of the cross-level slopes (95% confidence interval) to the error model and to sex differences among seasons. Additive: habitat type × sex + habitat type × season-year. Three-way: habitat type × sex × season-year. Per-point errors: Deming regression using the standard error of each estimate. Built-up areas were excluded at the landscape level in the per-point analyses. Bold values indicate confidence intervals that do not overlap zero.

| Habitat type | Additive (Table 2) | Additive, per-point errors | Three-way, per-point errors |
|---|---|---|---|
| Grasslands | **0.64 (0.24, 1.04)** | **0.62 (0.23, 1.00)** | **0.67 (0.01, 1.32)** |
| Winter cereals | 4.13 (−230.98, 239.23) | 4.89 (−65.44, 75.21) | 2.07 (−6.79, 10.92) |
| Fodder crops | **−1.67 (−3.24, −0.10)** | **−1.51 (−2.66, −0.35)** | −1.06 (−2.79, 0.67) |
| Other open fields | 3.55 (−15.48, 22.59) | 3.46 (−13.03, 19.95) | 4.00 (−9.08, 17.09) |
| Forests | **0.95 (0.65, 1.26)** | **1.08 (0.51, 1.64)** | 1.21 (−0.01, 2.44) |
| Forest edges | −3.88 (−8.18, 0.41) | −4.45 (−10.34, 1.44) | −5.24 (−21.01, 10.52) |
| Woodlands and hedgerows | −2.75 (−6.89, 1.39) | −3.26 (−9.00, 2.48) | −2.35 (−16.56, 11.87) |
| Farm tracks | **5.40 (3.52, 7.29)** | **5.18 (3.71, 6.65)** | **4.88 (1.20, 8.56)** |
| Secondary roads | 2.29 (−0.71, 5.28) | 2.65 (−1.32, 6.62) | 35.37 (−40.84, 111.59) |
| All habitats | 1.37 (−0.85, 3.58) | 2.57 (−0.38, 5.52) | 2.76 (−0.10, 5.63) |
